# A Bioluminescent Activity Dependent Platform, BLADe, for Converting Intracellular Activity to Photoreceptor Activation

**DOI:** 10.1101/2023.06.25.546469

**Authors:** Emmanuel L. Crespo, Akash Pal, Mansi Prakash, Alexander D. Silvagnoli, Zohair Zaidi, Manuel Gomez-Ramirez, Maya O. Tree, Nathan C. Shaner, Diane Lipscombe, Christopher I. Moore, Ute Hochgeschwender

**Affiliations:** College of Medicine, Central Michigan University, Mount Pleasant, MI 48859, USA; Biochemistry, Cell and Molecular Biology Graduate Program, Central Michigan University, Mount Pleasant, MI 48859, USA; Program in Neuroscience, Central Michigan University, Mount Pleasant, MI 48859, USA; Duke University, Undergraduate Neuroscience Program, Durham, NC 27710; Department of Neuroscience, Brown University, Providence, RI 02912, USA; University of California, San Diego, School of Medicine, Department of Neuroscience, 9500 Gilman Drive La Jolla, CA 92093-0662, USA; Carney Institute for Brain Science, Brown University, Providence, RI 02906, USA

## Abstract

Genetically encoded sensors and actuators have advanced the ability to observe and manipulate cellular activity, yet few non-invasive strategies enable cells to directly couple their intracellular states to user-defined outputs. We promote a bioluminescent activity-dependent (BLADe) platform that facilitates programmable feedback through genetically encoded light generation. Using calcium (Ca²⁺) flux as a model, we engineered a Ca²⁺-dependent luciferase that functions as an activity-gated light source capable of photoactivating light-sensing actuators. As an initial demonstration of the versatility of this platform we present two separate use cases in neurons. In the first application, the presence of luciferin triggers Ca²⁺ dependent local illumination that provides activity dependent gene expression by activating a light-sensitive transcription factor. In the second application, neuronal activity-driven Ca²⁺ fluctuations via locally generated bioluminescence control neural dynamics through opsin activation in single cells, populations and intact tissue. BLADe can be expanded to couple any signal that bioluminescent enzymes can be engineered to detect with the wide variety of photosensing actuators. This modular strategy of coupling an activity dependent light emitter to a light sensing actuator offers, in principle, a generalizable framework for state dependent cell-autonomous control across biological systems.

## Introduction

Bioluminescence, the emission of light generated when a luciferase enzyme oxidizes a small molecule luciferin, served as a non-invasive approach for *in vitro* and *in vivo* imaging for decades^1^. More recently, the development of brighter, blue light emitting luciferases in combination with light sensing elements, including opsins and non-opsin photoreceptors^2,3^, transformed bioluminescence from a mere imaging reagent to a versatile tool for controlling molecular processes. We and others have combined luciferases with channelrhodopsins and pumps for bioluminescence-driven changes in membrane potential^4–6^, with light-sensing transcription factors for gene expression^7–9^, with light-sensing enzymes for cAMP production^10^, as well as with genetically encoded photosensitizers for reactive oxygen species production^11,12^. The main advantages of using locally produced bioluminescence rather than external light sources for activating photoreceptors are the independence from optical fiber implants, thereby avoiding the physical damage to cells as well as from the limitation in the number of cells that can be simultaneously controlled.

A unique feature of genetically encoded light emitters is that they are amenable to molecular engineering. By inserting into the luciferase protein sensor moieties, light emission can be made dependent on the presence of various intracellular agents that report a cell’s state. Several such bioluminescent sensors have been developed for reporting intracellular biochemical signals, such as intracellular Ca²⁺, ATP, or cAMP, including for imaging studies *in vivo*^13^. Here, we are combining a bioluminescent sensor with light-dependent actuators to promote a general platform for converting biochemical signals within cells into photoreceptor activation and resulting downstream changes in cellular function. This Bioluminescence Activity Dependent (BLADe) paradigm serves to integrate cellular activity states with downstream consequences from transcription to membrane potential changes, depending on the optogenetic actuator coupled to the conditional luciferase. By engineering luciferases to respond to specific biochemical cues, this framework can, in principle, be extended to a broad range of intracellular states depending on the sensor domain used. Any bioluminescence indicator can, in principle, be transformed from a sensor into an activity integrator by using its light emission to activate photoreceptors.

As an initial demonstration of the versatility of this platform we employed a Ca²⁺ dependent luciferase. By making light production dependent on Ca²⁺ binding in a split luciferase construct, changes in Ca²⁺ levels are directly translated into bioluminescent light emission. Conversion of intracellular calcium level fluctuations into photoreceptor activation was tested in two separate applications in neurons. The Ca²⁺ dependent light emitter was paired with a light-dependent transcription factor that detected this light and affected transcription *in vitro* and *in vivo*. In another test case, when paired with channelrhodopsins Ca²⁺ dependent light emission enabled neural activity-driven change in firing activity *in vitro* in single neurons and neuronal populations; in mice, BLADe converted spontaneous activity and sensory evoked changes into activity dependent control of neocortical network dynamics.

## Results

### Engineering of a luciferase-based Ca²⁺ sensor

The *Gaussia princeps* luciferase variant sbGLuc is an efficient transmitter of photons/energy to opsin chromophores when tethered to opsins in luminopsins^5,14^. To make it Ca²⁺ dependent we split sbGLuc as was done previously for wildtype GLuc between Q88 and G89^15^ and introduced the Ca²⁺ sensing moiety CaM-M13 employing the M13 sequence from the Ca²⁺ sensor YC3.6^16^ and the calmodulin sequence from GCaMP6f^17^ (**Fig. 1a,b**). We generated a preliminary structural model that is, however, limited due to the lack of evolutionary structure data similar to the *Gaussia* luciferase protein structure (**Fig. 1c**). Nevertheless, we hypothesized that inserting the CaM-M13 domain between Q88 and G89 would allosterically control enzymatic activity by modulating the structural integrity of the native disulfide bond (C65–C77) which stabilizes the rigid α4–α7 helical core of sbGLuc (**Supplementary Fig. S1 and S2**). Assuming this mechanism, in the absence of Ca²⁺ the inserted calmodulin domain destabilizes this interaction, whereas Ca²⁺ binding promotes reassembly of the split halves and restoration of luciferase activity. We called this split sbGLuc based Ca²⁺ indicator Lumicampsin or LMC.

**Figure 1.**
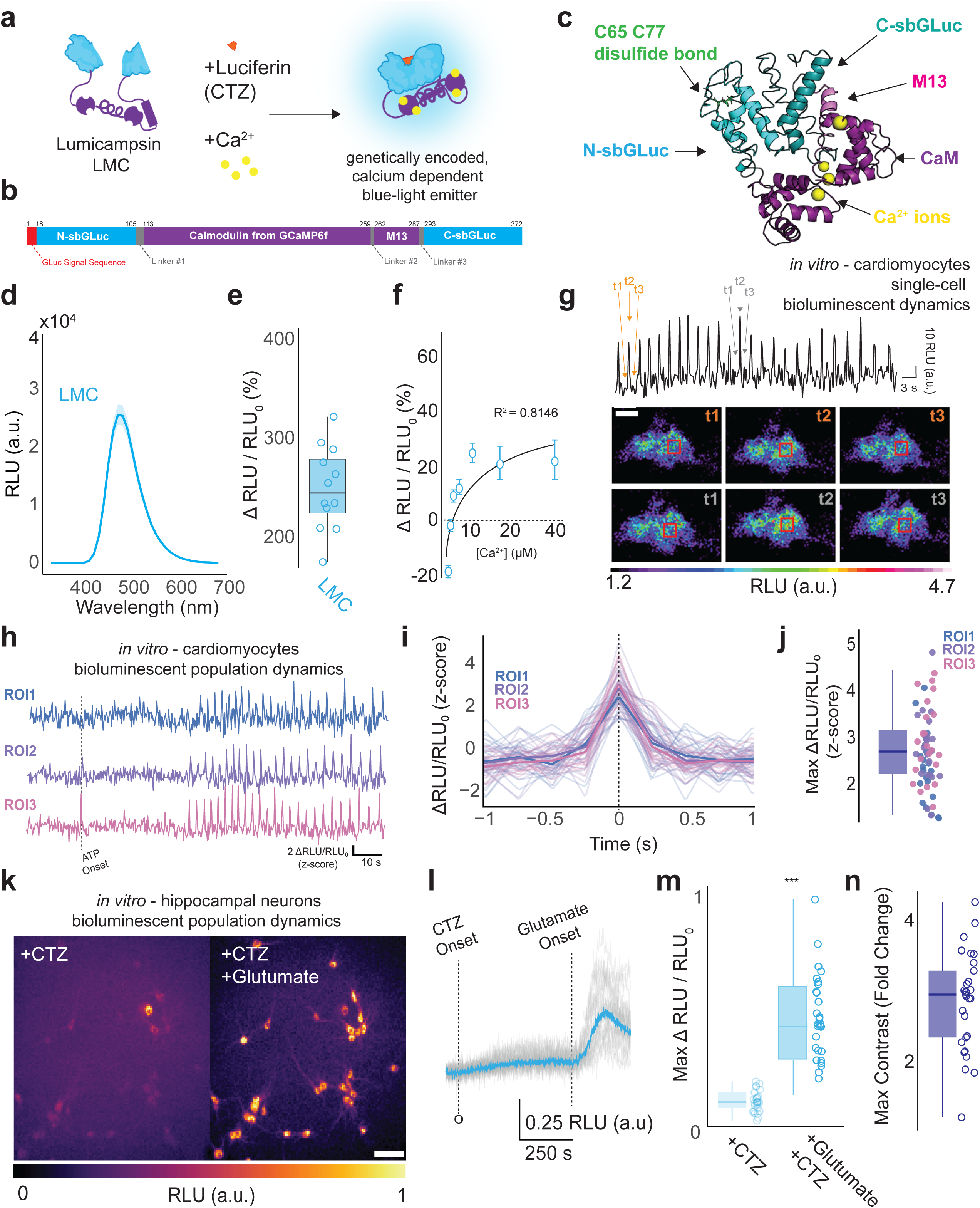
Design of a *Gaussia* luciferase based Ca^2+^ sensor. **a**. sbGLuc (light blue) is split by a CaM–M13 module (purple), producing a dim basal signal in the presence of its luciferin (CTZ). Binding of Ca²⁺ (yellow) reconstitutes an active luciferase and increases light emission in the presence of luciferin. **b**. Elements of the coding sequence for LMC: *Gaussia* luciferase signal peptide (red), N- and C-terminal fragments of sbGLuc (light blue), calmodulin sequence from GCaMP6f (purple), M13 sequence (purple), and various linker regions (gray). **c.** Structure model of LMC based on Alphafold2. **d-n.** LMC Ca^2+^ sensing in various contexts. **d, e.** HeLa cells. **d.** Emission wavelength for LMC. LMC construct was transiently expressed in HeLa cells and light emission after luciferin and histamine addition was measured across the wavelength spectra (data presented as mean ± SEM). **e.** Change in LMC luminescence in the presence of luciferin in response to 10 μΜ histamine in HeLa cells (N=12 wells). Boxplots show median, 25th and 75th percentiles (box edges), whiskers to most extreme data points, and individual outliers. **f.** HEK293 cells. Percent change in bioluminescence in HEK293 cells expressing LMC in response to 2 µM ionomycin under increasing extracellular Ca²⁺ concentrations. Cells were pre-incubated with 5 µM luciferin and varying concentrations of Ca²⁺; acute Ca²⁺ flux was initiated by injection of ionomycin before photon counting in a luminometer (mean ± s.e.m). **g - j.** Cardiomyocytes. **g.** Representative bioluminescence trace from an individual cardiomyocyte over time. Labels t1-t3 indicate troughs and peaks at two different time points (orange font, gray font). Pseudocolor images below show the same cell at the selected time points with ROI indicated by red box. Scale bar = 20 µm. **h.** Bioluminescence traces from regions of interest (ROIs) from three independent cardiomyocytes in culture (ROI3 represented in images in (g)), highlighting spontaneous and ATP-evoked activity. **i.** Individual bioluminescence transients from each ROI aligned to their peak and z-scored, with mean responses shown in bold. **j.** Quantification of peak z-scored bioluminescence responses across ROIs**. k - n.** Neurons. **k.** Representative hippocampal neurons expressing LMC imaged with luciferin alone (left) or with luciferin and glutamate (right). Scale bar = 100 µm. **l.** Mean (blue) overlaid with individual bioluminescence traces (grey) of LMC responses to bath-applied luciferin and glutamate. **m.** Box plot showing maximal LMC luminescence responses (Max ΔRLU/RLU₀) with luciferin alone vs glutamate plus luciferin (N= 29 neurons from two independent cultures, Wilcoxon signed rank test: p= 2.7023 ×10^−6^). **n.** Summarizes the maximal responses shown in (m), presented as the max fold change which was calculated per neuron as the ratio of peak responses in the CTZ+glutamate condition relative to the CTZ alone.

As the goal of LMC is to serve as a genetically encoded light source for activating blue light-sensing photoreceptors, we first confirmed its light emission spectra and Ca²⁺ responsiveness. HeLa cells were transfected with LMC, incubated with 5 µM luciferin (CTZ), and stimulated with histamine to induce a physiologically relevant increase in intracellular Ca²⁺. LMC retained a peak emission at roughly 490 nm (**Fig. 1d)** with a percent change in light emission before and after histamine addition of 232% (**Fig 1e**, IQR: 215–262%; range: 172–318%; N = 12 wells). Thus, LMC provides a blue-shifted optical report of intracellular calcium, enabling potential coupling to blue light sensitive optogenetic proteins required for the BLADe platform.

Experiments in HeLa cells confirmed that LMC responds to histamine-induced Ca²⁺ release at physiologic levels, but this system does not allow experimental control of intracellular Ca²⁺ levels in living cells. To address this, we transfected HEK293 cells with LMC and applied ionomycin under defined extracellular Ca²⁺ concentrations ranging from 0.4875 µM to 39 µM (**Fig. 1f**). LMC reported ionomycin-induced Ca²⁺ flux at all tested concentrations. The decrease in bioluminescent signal at lower extracellular Ca²⁺ conditions (0.4785µM -1µM) reflects the loss of intracellular Ca²⁺ as ionomycin moves Ca²⁺ from high (intracellular) to low (extracellular) concentrations.

To test the ability of LMC to report Ca²⁺ dynamics in cultured cells, we transfected primary cardiomyocytes with LMC and imaged them on DIV4 using an EMCCD camera. Cells were pre-incubated with luciferin (100 µM CTZ) for ∼10 min before imaging. Addition of 10 µM ATP increased contractility and induced rapid Ca²⁺ transients with robust peaks (**Fig. 1g-j**).

Next, to test whether LMC could report neurotransmitter-evoked Ca²⁺ influx in neurons, we transfected primary hippocampal cultures and perfused luciferin (CTZ) followed by luciferin with glutamate. Bioluminescent signals were low with CTZ alone but increased after glutamate application **(Fig. 1k**). Time-locked traces showed a clear rise in bioluminescence following glutamate onset (**Fig. 1l**). Across individual neurons, we observed an increase in normalized peak responses with glutamate compared to luciferin alone (**Fig. 1m**). In addition, responses also exhibited high contrast relative to baseline, with a median fold-change of ∼2–4 across neurons (**Fig. 1n**).

### Bioluminescence as an activity-gated light source

We next compared LMC with other published bioluminescent Ca²⁺ indicators to evaluate its baseline emission and suitability as an activity-gated light source. HeLa cells were transiently transfected with the *Gaussia* luciferase based LMC, and the *Oplophorus* luciferase based BlueCaMBI^18^, GLICO^19^, or GeNL^20^ and light emission was assayed in a luminometer following luciferin addition (CTZ for LMC, hCTZ for the other three sensors) and histamine stimulation. Time-series traces show that BlueCaMBI, GLICO, and GeNL, produced higher overall photon output than LMC (**Fig. 2a**). Although these sensors were brighter, they also exhibited substantial baseline photon emission in the presence of luciferin under resting Ca²⁺ conditions, relative to their response after histamine stimulation (**Fig. 2b**). For imaging Ca²⁺ fluctuations the baseline is defined as the value immediately before histamine addition, and Ca²⁺ spikes are represented by these before/after values. However, bioluminescent sensors will not be suitable as drivers of photoreceptors unless their light emission in the initial presence of luciferin at baseline is below the activation limit of photoreceptor activity. All three indicators—BlueCaMBI, GLICO, and GeNL—have been reported to capture Ca²⁺ fluctuations, but in those studies, bioluminescence signal traces were typically normalized following luciferin pre-incubation^18–20^. While this normalization approach is appropriate for imaging relative Ca²⁺ dynamics, the high absolute photon emission at resting levels of Ca²⁺ limits their use as activity-gated light sources required for BLADe activation of photoreceptors.

**Figure 2.**
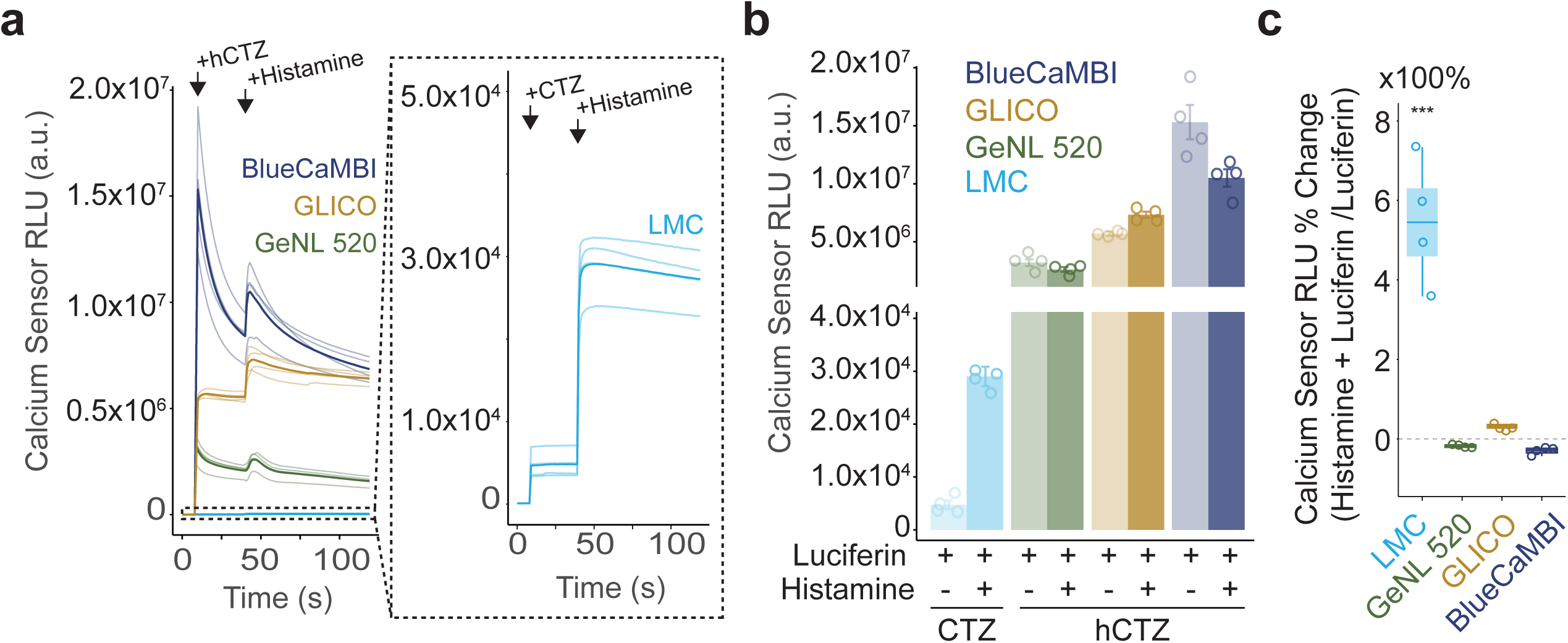
Integrating Ca^2+^ flux with activation of photoreceptors. **a.** Luminometer time series from HeLa cells transiently expressing BlueCaMBI, GLICO, GeNL, or LMC following addition of luciferin and histamine. Inset shows LMC traces on an extended scale. **b.** Peak bioluminescence measured after luciferin and histamine stimulation for each sensor from (a). **c.** Histamine-specific percent change in bioluminescence normalized to luciferin baseline, expressed as percent change for each calcium sensor. (N=4 wells/sensor, LMC vs GLICO: *P=* 4.65×10^−6^, GeNL: *P=* 5.51×10^−7^, BlueCAMBI: *P=* 1.46×10^−6^; GeNL vs GLICO *P*= 0.38, BlueCAMBI P= 0.996; BlueCAMBI vs GLICO: *P*=0.723, Tukey’s post hoc multiple-comparison’s test following a one-way analysis of variance (ANOVA), F(_3, 12_) **=** 48.56, *P**=*** 5.51×10^−7^). Boxplots show median, 25th and 75th percentiles (box edges), whiskers to most extreme data points, and individual outliers.

High absolute brightness benefits imaging, but BLADe optogenetic applications require low basal emission and a large absolute increase in light output during Ca²⁺ influx. When a sensor is used for coupling light emission with photoreceptor activation, the critical baseline value is the level of light emission once the luciferin enters the cell. If that initial level is too high, the photoreceptor will be activated at baseline, rather than being activated as a consequence of increased levels of Ca²⁺. Consistent with this, when normalized to the luciferin baseline, only LMC exhibited a large relative change in photon emission following histamine-induced Ca²⁺ release (**Fig. 2c**). Accordingly, despite lower absolute brightness, the unique low background and high dynamic range of LMC positions it as an activity-gated light source ideally suited for converting intracellular signals into optical activation of downstream photoreceptors. At resting calcium levels, we hypothesize LMC light emission remains dim and is below the threshold needed to activate photoreceptor chromophores, minimizing spurious activation of light sensing proteins. We next asked whether this activity-dependent light source could drive optogenetic tools as the basis of the BLADe platform.

### Integrating Ca²⁺ flux with change in transcription

LMC expressing cells showed low baseline bioluminescence in the presence of luciferin and a strong increase in light output following Ca²⁺ influx. We reasoned that LMC’s activity-gated light emission could be used to photoactivate light-sensitive transcriptional effectors. To test this prediction, we co-expressed LMC with the light sensing transcription factor EL222^21^. In the presence of blue light, EL222 dimerizes and then binds to its promoter 5xC120, leading to transcription of the reporter gene (**Fig. 3a**).

**Figure 3.**
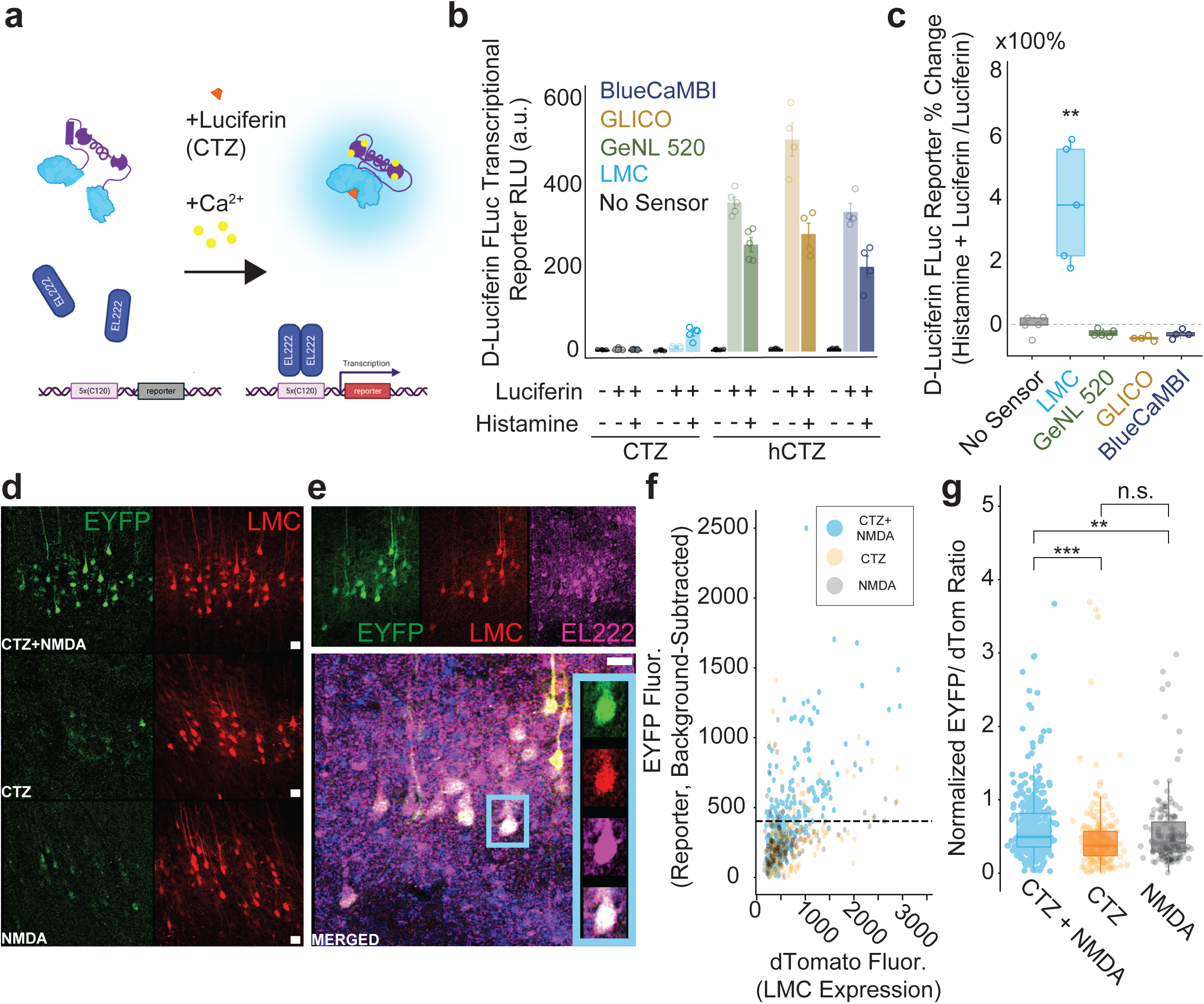
Converting intracellular activity to transcription. **a.** Schematic of LMC leading to bioluminescence mediated dimerization of the light sensing transcription factor EL222 in the presence of Ca^2+^ and luciferin (CTZ). The photoactivated EL222 homodimer then binds to the promoter 5xC120 and leads to transcription of the reporter gene. **b.** Firefly luciferase (FLuc) transcriptional reporter measured following no treatment (no luciferin, no histamine), luciferin-only, or luciferin and histamine treatments in HeLa cells transiently expressing BlueCaMBI, GLICO, GeNL, or LMC. **c.** FLuc transcriptional reporter normalized to luciferin baseline expressed as percent change for each sensor/condition. N = 4-5 wells per sensor; LMC vs BlueCaMBI: P = 0.00998; vs GLICO: P = 0.00998; vs GeNL 520: P = 0.00609; vs no calcium sensor: P = 0.00609; one-sided Wilcoxon rank-sum tests following a Kruskal–Wallis (non-parametric ANOVA) test across sensors: H = 16.1, df = 4, P = 0.00287. **d.** Representative confocal images showing EYFP transcriptional reporter expression (green) in mouse prefrontal cortex following NMDA alone, luciferin (CTZ) alone, or combined CTZ and NMDA administration. Red fluorescence marks neurons expressing the LMC-P2A-dTomato construct. Scale bar = 100 µm. **e.** Individual channels are shown separately with a merged image below. Insets highlight representative neuron demonstrating co-localization of EYFP (green), LMC (red), and EL222 (magenta) expression. Scale bar = 50 µm. Merged image includes DAPI and scale bar = 20 µm. **f.** EYFP reporter expression quantified as per-cell EYFP/dTomato ratio across treatment groups. Each point represents a single cell derived from three treatment groups: Luciferin-NMDA (N= 350 cells/2 mice), Luciferin-ONLY N= 197 cells/2 mice), NMDA-ONLY (N= 147 cells/2 mice). Dotted line indicates the 90th percentile EYFP/dTomato ratio observed in the NMDA-ONLY group, used as a threshold to define transcriptional activation above baseline. **g.** EYFP reporter intensity plotted against dTomato fluorescence for each cell and condition. Boxplots show median, 25th and 75th percentiles (box edges), whiskers to most extreme data points, and individual outliers.

HeLa cells expressing EL222, the Firefly luciferase (FLuc) reporter, and each calcium sensor were either left untreated, treated with luciferin for 15 minutes, or treated with luciferin and histamine for 15 minutes. Transcription of the FLuc reporter was measured 18 hours later as luminescence after addition of D-luciferin to the cells (**Fig. 3b**). Left untreated or in the absence of a sensor, no EL222 activation was observed. With addition of luciferin (CTZ for LMC, hCTZ for the other sensors) transcription at baseline Ca²⁺ levels is very low for unreconstituted LMC, in contrast to the other three sensors. The addition of luciferin and histamine should reflect the increased light emission of the reconstituted – due to the histamine-induced increase in intracellular Ca²⁺ – luciferase resulting in higher EL222 activation and higher transcription of FLuc. This prediction is seen with LMC, but not with the other three Ca²⁺ sensors, due to their high baseline signal. Interestingly, the absolute FLuc reporter expression for BlueCaMBI, GLICO, and GeNL is lower in cells exposed to luciferin and histamine than in cells exposed to luciferin alone. This is likely a reflection of the increased consumption of the luciferin in the presence of increased Ca²⁺, resulting in decreasing the time of light emission for EL222 activation.

All calcium dependent indicators produced FLuc transcription under luciferin treatment alone, indicating that transcription can be photoactivated even at very low levels of calcium. In the presence of luciferin, baseline transcription was strongest for BlueCaMBI, GLICO, and GeNL, which were markedly brighter than LMC — a trend consistent with their higher baseline bioluminescence in luminometer traces (**Fig. 2b**). In contrast, only LMC showed a calcium-dependent increase in transcription, with a significant increase in FLuc signal in the luciferin + histamine condition relative to all other sensors (**Fig. 3c**). LMC maintains a low luciferin baseline and undergoes a discrete calcium-dependent increase in light output, preserving a clear calcium-state transition. Moreover, the overall transcriptional output from LMC remained modest compared to the other sensors, which is consistent with its lower total light output in luminometer assays (**Fig. 2b**). As a result, effective photoreceptor gating in the BLADe system depends not on absolute light output but on preservation of a discrete calcium-state transition above a low luciferin baseline. Having established that LMC produces calcium-dependent and - state-specific light sufficient to activate EL222 *in vitro*, we next tested whether the BLADe transcriptional system could function *in vivo*.

As a pilot study, we tested whether LMC could drive transcription in neurons *in vivo* when network activity was chemically elevated with NMDA. We performed stereotactic injections of AAV9-hSyn-LMC-P2A-dTomato, AAV9-CAG-EL222, and AAV9-5×C120-EYFP into the prefrontal cortex of anesthetized mice. After virus delivery, the craniotomy was immediately covered to further minimize unintended light exposure during recovery. After 2–3 weeks, to allow for maximal expression, mice received intraventricular injections of either NMDA + vehicle, luciferin (CTZ) alone, or luciferin + NMDA. These conditions allowed us to test: (1) non-specific background transcriptional activity in the absence of light (NMDA + vehicle), (2) light-dependent transcription in the context of spontaneous activity (luciferin alone), and (3) BLADe-mediated transcription (luciferin + NMDA).

Confocal imaging of tissue collected ∼20 hours after treatment showed increased EYFP reporter labeling in individual LMC-expressing neurons from mice treated with luciferin + NMDA, compared to the other treatment groups (**Fig. 3d,e**). Single-cell analysis confirmed that EYFP reporter increase occurred across a range of dTomato expression levels (**Fig. 3f**), indicating that the Ca²⁺-dependent enhancement of transcription was not due to differences in viral expression of LMC. Quantification of EYFP reporter/dTomato tool expression ratios revealed higher reporter expression in the luciferin + NMDA group compared to both the luciferin-only group and the NMDA-only group (**Fig. 3g**, Wilcoxon rank-sum test, Bonferroni-corrected luciferin+NMDA vs luciferin: *P*= 1.62 × 10⁻⁷; luciferin+NMDA vs NMDA: *P*= 0.0057; luciferin vs NMDA: *P* = 0.196; pooled from FOVs of 197-350 cells/N=2 mice for each treatment group). The luciferin-only and NMDA-only groups were not significantly different from each other suggesting that reporter activation requires both intracellular calcium and bioluminescent light, as required with LMC-BLADe gating. Together, these findings are consistent with BLADe functioning *in vivo* in the mouse brain to drive calcium-dependent transcription, with the overall magnitude of reporter labeling remaining modest. Increasing light output of the Ca²⁺ dependent luciferase, improving the contrast between active and inactive cells, and simplifying viral expression strategies may optimize this system for robust transcriptional labeling in future applications.

### Integrating Ca²⁺ flux with change in membrane potential

To demonstrate the modularity of the BLADe platform, we tested whether LMC could drive light-sensitive ion channels. Following an action potential, the influx of intracellular Ca²⁺ provides a convenient and high-resolution proxy for BLADe neural activation. We reasoned that LMC could detect these calcium transients and, through co-expression with an excitatory opsin, rapidly alter spiking dynamics in an activity-dependent manner. To this end, we applied step current injections to hippocampal neurons *in vitro* in the presence of synaptic blockers to isolate cell-intrinsic responses. We then performed whole-cell patch-clamp recordings on neurons co-transformed with LMC and an opsin (**Fig. 4a**). By visualizing co-localized expression of an opsin-EYFP (green; here ChR2(C128S)) and the Ca²⁺-dependent luciferase LMC-P2A-dTomato (red), we selected these double-positive neurons for testing the integration of cellular states via BLADe opsin activation.

**Figure 4.**
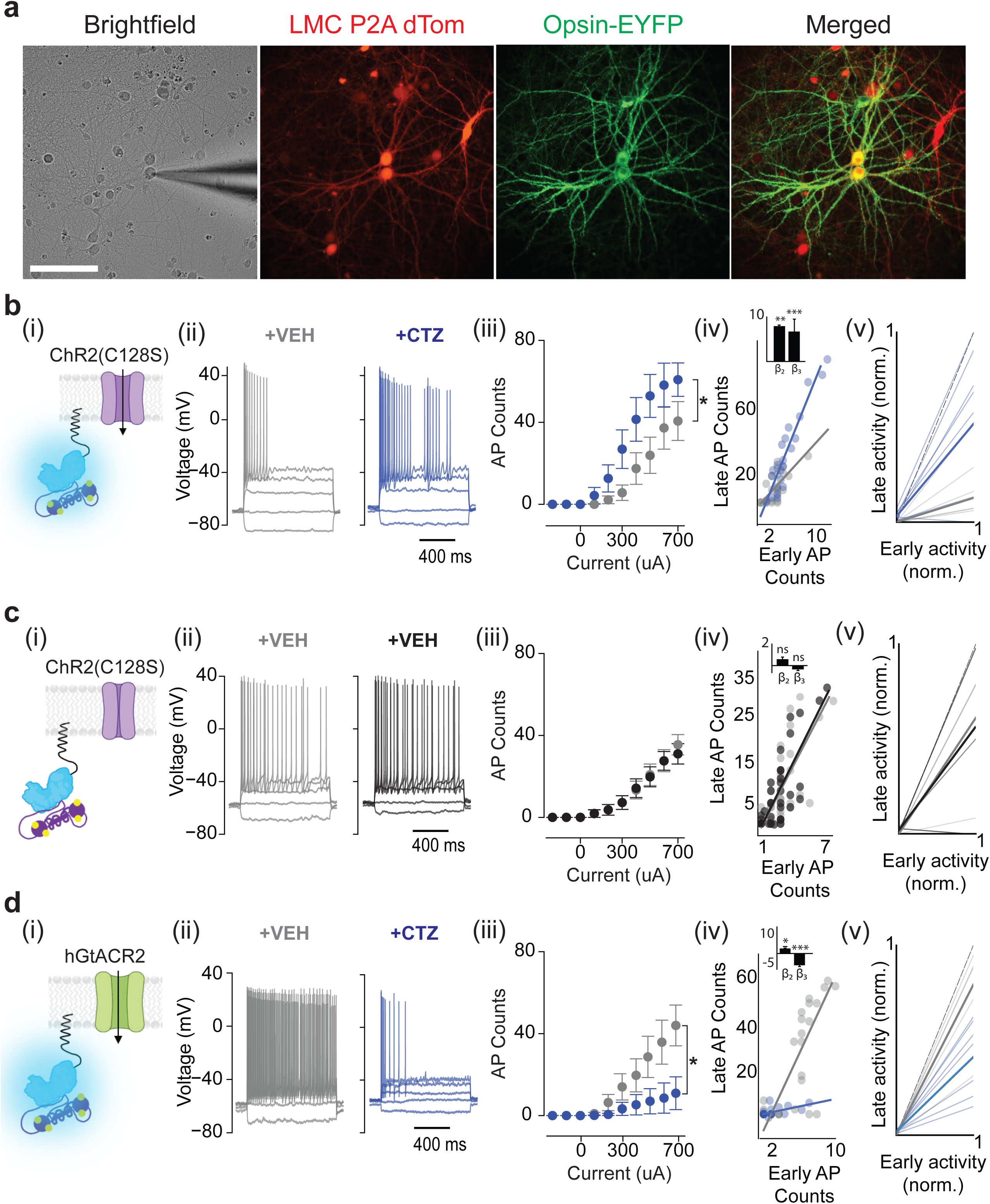
Integrating Ca^2+^ flux with change in membrane potential in single neurons *in vitro*. **a.** Brightfield and fluorescence images of cultured cortical neurons nucleofected with opsin (ChR2(C128S))-EYFP and LMC-P2A-dTomato and used for whole-cell recordings. The patch electrode is visible in the brightfield image. Scale bar = 50 µm. **b.** (i) Schematic of farnesylated LMC anchored to the inner cell membrane near the excitatory opsin ChR2(C128S) enabling light-mediated activation in the presence of depolarization-induced Ca²⁺ flux and the luciferin (CTZ). (ii) Representative membrane voltage responses to step current injections before (gray) and after (blue) bath perfusion of luciferin in a neuron co-expressing LMC and ChR2(C128S). (iii) Action potential counts under vehicle (gray) and luciferin (blue) conditions. Data expressed as mean ± SEM. (iv) Trial-level regression relating late firing to early firing across sweeps. (v) Normalized across-neuron early–late lines showing per-cell transformations before and during luciferin. **c.** (i) Schematic of a control condition, where LMC is co-expressed with ChR2(C128S) in the presence of Ca²⁺ but without luciferin, preventing light-mediated activation. (ii–v) Same analyses as in b. **d.** (i) Schematic of LMC co-expressed with the inhibitory opsin hGtACR2, enabling light-mediated suppression of activity in the presence of Ca²⁺ and luciferin. (ii–v) Same analyses as in b, showing inhibition rather than activation. Data are presented as mean ± SEM in panels.

We predicted that LMC generates light directly in response to Ca²⁺ influx which occurs during action potential generation induced by depolarizing current injections. We reasoned that increasing levels of depolarizing current in the presence of the luciferin would produce enough bioluminescence to further photoactivate ChR2(C128S)^22^ via a cell-autonomous, positive feedback optical loop. In line with our prediction, luciferin perfusion during the same depolarizing current elicited significantly more detected action potentials than during vehicle perfusions **(Fig 4b, ii and iii,** N=9 neurons, p= 4.954×10^−9^, paired sample Wilcoxon signed rank tests), indicating a light-dependent, cell-autonomous control of membrane voltage potential.

These initial findings are consistent with LMC bioluminescence being sufficient to photoactivate the opsin, but they do not directly prove BLADe control or reveal how the effect emerges over the course of the 1-second depolarizing step. Given that LMC reports spontaneous intracellular calcium dynamics with sub-second temporal resolution (**Fig. 1**), we hypothesized that any optical feedback effect should manifest within the timescale of the depolarizing step. If BLADe control depends on activity-coupled light production, then the timing of spiking should reflect this coupling, i.e. the effect of LMC-driven opsin activation should evolve across the depolarizing step. To this end, we first measured how spikes were distributed across the 1-second depolarizing step using a normalized early–late comparison per trial to detect timing shifts rather than changes in overall firing (see Methods). The spike-timing distribution shifted toward the end of the 1-second depolarizing current during luciferin perfusion in neurons co-expressing LMC and ChR2(C128S), in an opsin- and CTZ- dependent manner (**Supplementary Fig. S3**). Our observations support a model in which LMC-driven bioluminescence engages the opsin through a voltage-dependent feedback process and prompted us to characterize how LMC alters the relationship between early and late spiking across the depolarizing stimulation.

We reasoned LMC-driven bioluminescence is not merely a tonic light source but may also act as an activity-coupled light emitter, such that its optogenetic effect on neuronal firing scales with the level of spiking activity within a trial. We therefore divided each trial into an early window (0–200 ms following stimulation onset) and a late window (>200 ms), capturing initial versus sustained spiking activity under continued current stimulation. To separate tonic light effects from activity dependent modulation and quantify how LMC integrates ongoing spiking activity into later responses, we applied a trial-level linear model relating late firing to early activity, luciferin condition, and their interaction, pooled across neurons (see Methods). In this model, β₁ captures the relationship between early and late phase spiking in the vehicle condition, reflecting the baseline relationship between early and late spiking within a trial. In contrast, the main luciferin effect term (β₂) reflects an activity independent shift in late firing, consistent with tonic optogenetic drive. The interaction term (β₃) captures activity dependent modulation (gain), reflecting amplification (or suppression) of spiking that scales with ongoing activity.

Accordingly, a tonic optogenetic effect in the presence of luciferin would manifest as a uniform shift in firing (β₂) with the direction of the shift (additive: β₂ > 0; subtractive: β₂ < 0) determined by the biophysical properties of the co-expressed opsin. In contrast, activity-dependent modulation would be expressed as a change in the slope linking early to late activity (β₃ ≠ 0). Thus, we predicted that LMC-driven bioluminescence during luciferin perfusion would rapidly shape early–late epoch spike coupling, with positive β₃ values indicating multiplicative gain with excitatory opsins and negative β₃ values indicating divisive suppression with inhibitory opsins.

For neurons expressing the excitatory opsin (**Fig. 4b iv**), luciferin increased the slope relating early to late phase spiking, indicating that early epoch activity more strongly predicted late responses and that this relationship was multiplicatively scaled across trials (β_1_= 3.927, CI [1.49, 6.37], p= 0.002, β_2_= 9.33, CI [3.63, 15.02] p= 0.002, β_3_= 5.02, CI [2.30, 7.74], p = 0.001). Across recordings, multiplicative gain was observed in most neurons (**Fig. 4b v**), with 6/9 recordings exhibiting increased slope relative to baseline and the remainder showing either no change (1/9) or reduced slope (2/9). Together, in the presence of luciferin, LMC converts early spiking activity into activity-dependent gain, amplifying late responses in proportion to the magnitude of spiking activity established during the onset of stimulation.

In contrast, neurons co-expressing LMC and ChR2(C128S) that received two sequential vehicle perfusions showed no changes in spiking dynamics. Overall firing rate and the temporal distribution of spikes across the stimulus window was unaltered across trials **(Fig. 4c ii-iii**, N=14 neurons, *p=* 0.3117, paired sample Wilcoxon signed rank test). Consistent with this observation, the linear model revealed no evidence of either tonic modulation or activity dependent changes in gain (**Fig. 4c iv**, β_1_= 4.54, CI [3.29, 5.78], p= 0.001, β_2_= 0.55, 95% CI [-2.39, 3.49] p= 0.71, β_3_= 0.23, CI [-1.54, 2.00], p= 0.794). No consistent change in slope was observed, with most recordings (10/14) showing no measurable change and a subset of neurons (4/14) exhibiting reduced slopes (**Fig. 4c v**). Our result confirms that the observed bias in spike timing requires luciferin, and by extension, LMC stimulus-dependent bioluminescence.

To confirm these changes depend on a functional opsin rather than bioluminescence alone, we next tested a non-photoconductive ChR2 mutant, ChR2(C128S)-E97R-D253A^23,24^ (“DUD” opsin) alongside LMC (**Supplementary Fig. S4**). There were no observed luciferin-induced shifts in spike timing or firing rate, indicating that BLADe control does not alter the intrinsic excitability or temporal dynamics of spike emergence across trials in the absence of a functional opsin.

Building on our findings with the excitatory opsin, we next tested whether LMC could suppress firing in neurons expressing an inhibitory opsin. The same activity-coupled mechanism is expected to produce divisive suppression, such that increases in early spiking reduce subsequent firing later in the stimulus epoch. In line with our prediction, neurons co-expressing hGtACR2^25^ and LMC exhibited fewer total spikes (**Fig. 4d, ii and iii**, N=10 neurons, p*=.* 0.3117, paired sample Wilcoxon signed rank test) and weakened the relationship between early and late spiking after accounting for tonic optogenetic effects (**Fig. 4d, iv,** β_1_= 7.22, CI [5.36, 9.07], p= 0.001, β_2_= -11.44, CI [-20.89, -2.003], p=0.019, β_3_= -6.489, CI [-20.89, -2.003], p= 0.019). Across recordings, divisive suppression was observed in most neurons (**Fig. 4d v**), with 7/10 recordings exhibiting reduced slope relative to baseline and the remainder showed no measurable slope change (3/10).

Moving beyond single-cell recordings, we next tested how LMC modulates firing rates at the network level by performing extracellular recordings on cultured cortical neurons plated on multi-electrode arrays (MEA)^26^. We virally co-transduced neurons with an excitatory opsin ChR2(C128S)-EYFP and the Ca²⁺-dependent luciferase LMC-P2A-dTomato (**Fig. 5a,b**). Time-locked multi-unit activity (MUA) traces revealed rapid population-level activity following luciferin or vehicle application (**Fig. 5c**). While vehicle addition alone can transiently increase network activity in MEAs—likely due to mechanical or ionic effects of media perfusion—these responses were brief and stereotyped across independent experiments. To enable comparison, we aligned all trials to the application artifact, allowing time-locked analysis between vehicle and luciferin treatments. In doing so, we could distinguish true activity-dependent changes from non-specific network effects caused by vehicle addition. Vehicle and luciferin treatments were randomized on the same MEA culture. All comparisons used the same electrode sites with recording ∼4-5 hours apart.

**Figure 5.**
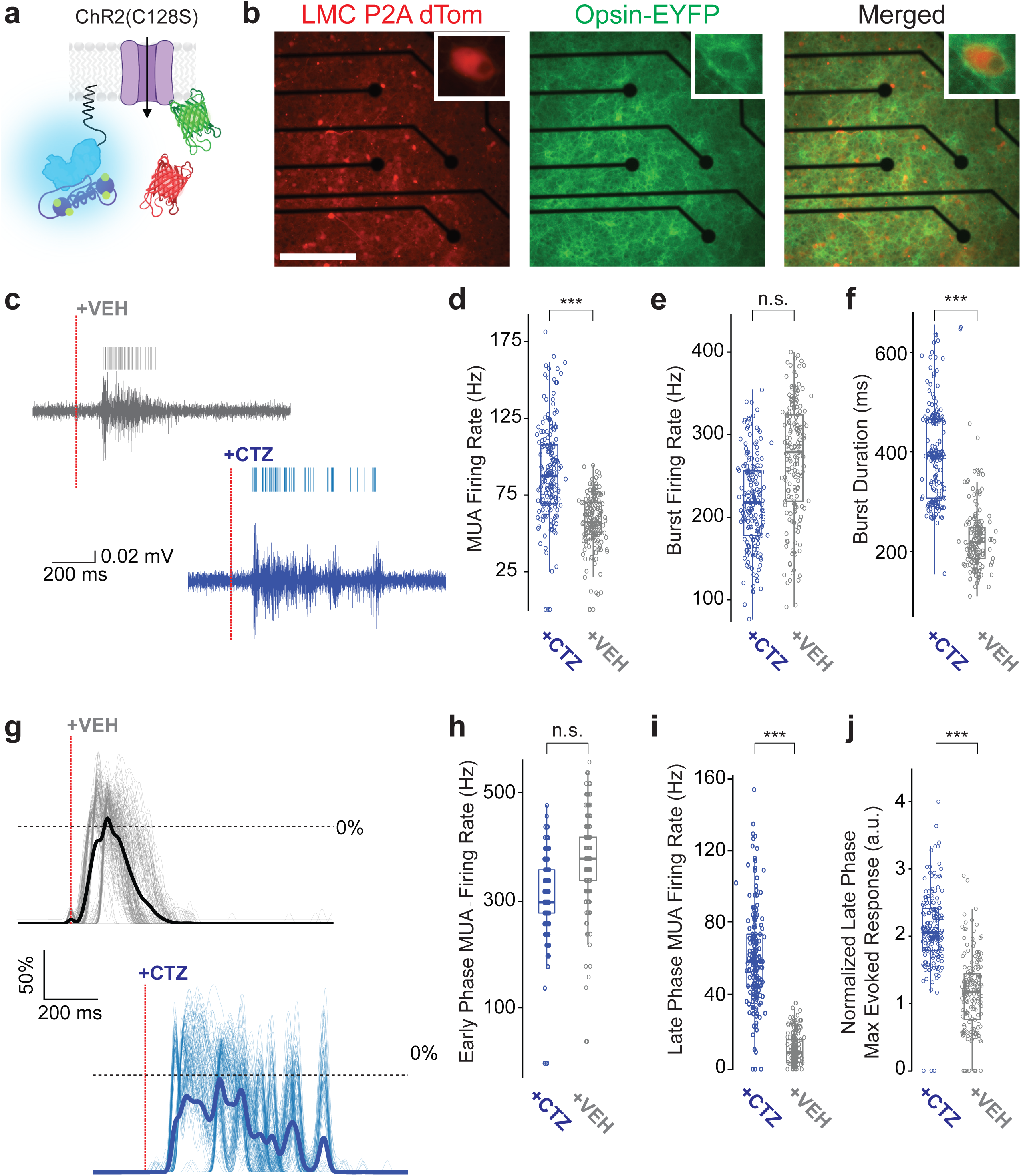
Integrating Ca^2+^+ flux with change in membrane potential in neural populations *in vitro*. **a.** Schematic of farnesylated LMC anchored to the inner cell membrane near the excitatory opsin ChR2(C128S) enabling light-mediated activation in the presence of depolarization-induced Ca²⁺ flux and the luciferin (CTZ). **b.** Fluorescence images of cultured cortical neurons co-transduced with LMC-P2A-dTomato (red) and ChR2(C128S)-EYFP (green). **c.** Representative filtered voltage traces from one channel before and after vehicle (top) or luciferin (bottom) application. Red dashed line indicates time of application. **d.** MUA firing rate was increased in the luciferin condition (Wilcoxon signed-rank test, *P*= 3.93 × 10⁻²⁸, Holm-corrected). **e.** Burst rate was not significantly different between luciferin and vehicle conditions (Wilcoxon signed-rank test, *P*= 0.63, Holm-corrected). **f.** Burst duration was significantly prolonged in the luciferin condition (Wilcoxon signed-rank test, *P*= 3.18 × 10⁻⁴⁹, Holm-corrected). **g.** Time-aligned MUA traces across all electrodes showing normalized population responses following vehicle (top) or luciferin (bottom) application. Individual electrode traces are shown in grey (vehicle) and blue (luciferin); thick lines represent the mean response per condition. MUA was smoothed using a sliding 5-point boxcar window. **h.** Early-phase (0–50 ms) MUA firing rates following application was not different between luciferin and vehicle (Wilcoxon signed-rank test, *P*= 0.13, Holm-corrected). **i.** Late-phase (150–1000 ms) MUA firing rate was significantly higher in the luciferin condition (Wilcoxon signed-rank test, *P*= 2.61 × 10⁻⁸, Holm-corrected). **j.** Late phase normalized MUA was significantly increased under luciferin (Wilcoxon signed-rank test, *P*= 1.02 × 10⁻¹¹, Holm-corrected). For **d-f** and **h-j**, N=57-59 pairs of electrodes per recording pooled across three independent MEA cultures. Boxplots show median, 25th and 75th percentiles (box edges), whiskers to most extreme data points, and individual outliers.

In the presence of luciferin, MUA was enhanced relative to vehicle one second after application (**Fig. 5d**). Burst firing rates were similar between conditions (**Fig. 5e**), yet burst duration was significantly longer with luciferin (**Fig. 5f**). With burst firing rates unchanged but MUA elevated, we next tested whether neural activity was prolonged over time due to co-expression of an excitatory opsin with LMC. To this end, we analyzed the early phase as defined by the 0–50 ms window beginning immediately after the application artifact, corresponding to the initial rise in MUA relative to baseline (**Fig. 5g**). The time-locked window allowed for reliable comparisons of immediate firing between treatment conditions. The late phase was defined as the remaining portion of the 1-second window, from 100–1000 ms post-application, allowing us to test whether LMC modulates both the onset and sustained components of network activity.

Early phase firing rates were similar between vehicle and luciferin conditions indicating that initial responses to perfusion were comparable across treatments (**Fig. 5h**). Thus, the observed effects are not due to media-driven excitability or non-specific stimulation. In contrast, the late-phase response was significantly enhanced in the presence of luciferin (**Fig. 5i).** When normalizing the late-phase MUA to the early-phase, we observed a clear enhancement in MUA consistent with LMC activation with luciferin (**Fig. 5j**). The observed sustained effect on network excitability in the presence of the luciferin suggests that LMC-driven light production prolongs neuronal activity via local activation of the co-expressed excitatory opsin. Finally, to test whether LMC can bidirectionally modulate population activity, we co-expressed LMC with the inhibitory opsin hGtACR2 and observed luciferin-dependent suppression of MUA, confirming that inhibition requires LMC expression (**Supplementary Fig. S5,** median ΔFR = +7 Hz, *n* = 19 vs median ΔFR = –14 Hz, *n* = 51 with LMC, Wilcoxon rank-sum test, p= 1.576 × 10^−10^). Our MEA experiments extend our single-cell findings by testing whether LMC optogenetically modulates neural activity at the population level in cultured neurons. However, because spiking dynamics are not resolved on a trial-by-trial basis in this preparation, we next tested whether LMC mediates activity-dependent gain *in vivo* using extracellular recordings.

Building on our *in vitro* electrophysiology observations, we next wanted to determine if LMC can integrate physiological sensory inputs to alter neocortical networks. We leveraged the well-characterized vibrissae circuit to ask whether LMC can detect sensory cues and reshape how they are encoded within the neocortex. To this end, we injected Cre-dependent versions of LMC and the excitatory opsin CheRiff^27^ into the left cortex of Emx1-Cre mice to target both constructs to excitatory pyramidal neurons within the vibrissa region of primary somatosensory cortex (vSI). Three to four weeks later, we then performed extracellular recordings with laminar silicon probes in vSI. As a control, a separate cohort of Emx1-Cre mice received identical surgical procedures and recordings but did not express the opsin (N = 2 mice per group). During recordings in head fixed anesthetized mice, we perfused ACSF or ACSF containing luciferin (CTZ) over the cortical surface while delivering vibrissal deflections (**Fig. 6a, b**).

**Figure 6.**
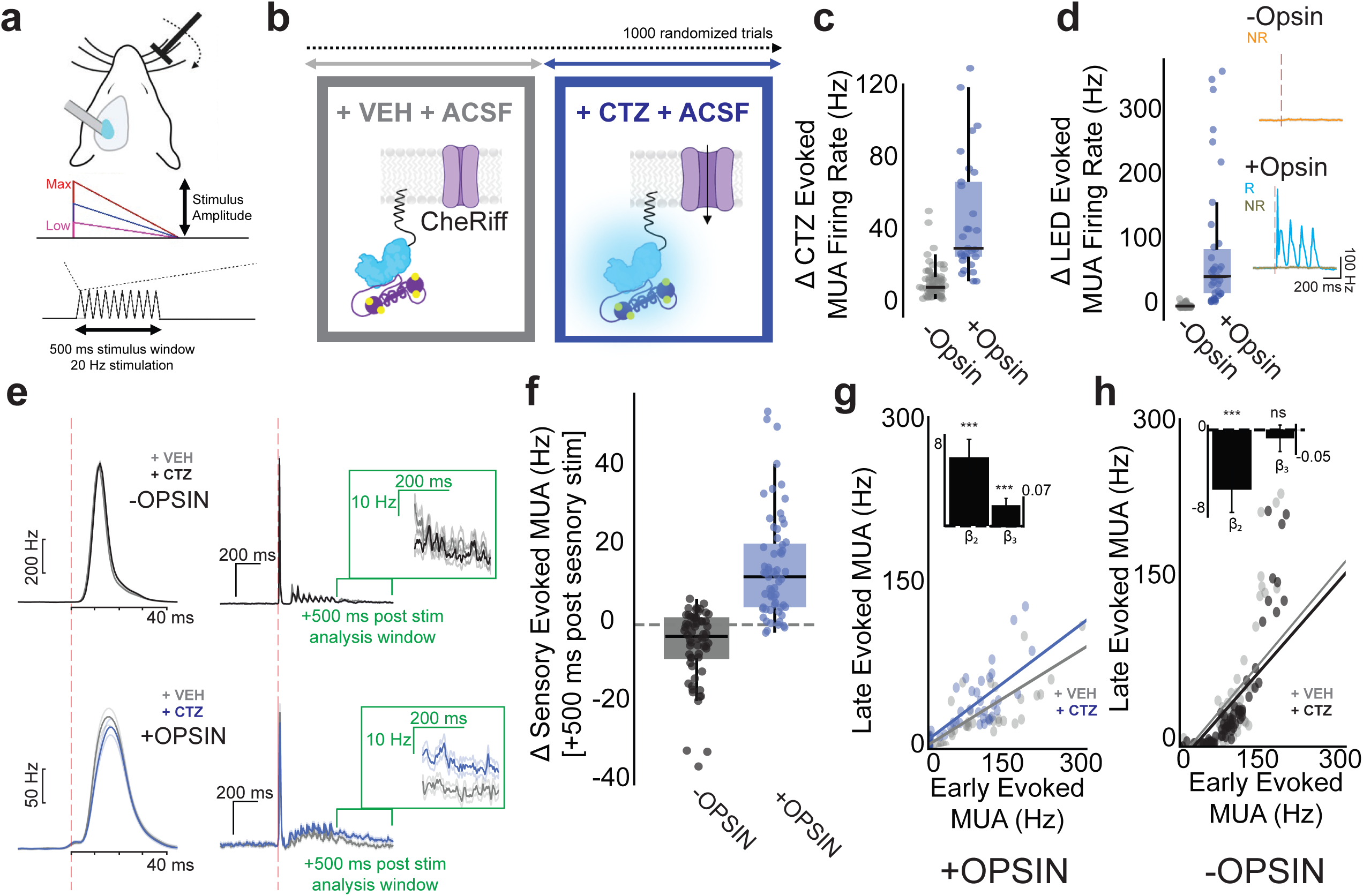
Integrating spontaneous and sensory inputs into changes in firing rates *in vivo*. **a**. Schematic of the experimental design. Extracellular electrophysiology recordings from SI barrel cortex during varied vibrissal deflections (zero, low, mid, max intensities) for 500ms at 20Hz. **b**. Schematic of the experimental timeline. Each recording consisted of 1000 trials (∼2.5 hour sessions) where ACSF containing vehicle or CTZ was constantly perfused over the craniotomy. During each treatment epoch, vibrissal deflections were randomly presented. In the presence of the sensory stimulus and vehicle (grey box), there is no activity dependent light emission of LMC. In contrast, in the presence of CTZ and the sensory stimulus (blue box), sensory recruitment leads to activity dependent bioluminescence and subsequently photoactivation of opsins in neurons. **c**. Luciferin (CTZ) application increases spontaneous multiunit activity (MUA) in opsin-expressing mice relative to no-opsin controls. **d.** LED-evoked MUA responses in opsin-expressing mice exceed the 95th percentile of the no-opsin null distribution. *Inset:* MUA traces from a non-responsive channel (NR; orange) in a −opsin cohort and from responsive (R; teal) and non-responsive (NR; green) channels in a +opsin cohort. The red dashed line indicates LED onset. **e.** Sensory-evoked responses in vibrissae S1 from LMC-expressing mice (N = 2 mice per group). *Left*, mean ± SEM baseline-corrected multiunit activity (MUA) responses to a single whisker deflection during vehicle (grey) and after CTZ perfusion (blue) in opsin-expressing mice; corresponding responses from no-opsin control mice are shown in grey (vehicle) and black (CTZ). *Middle*, peristimulus time histogram (PSTH) of MUA during the full whisker stimulation epoch (1500 ms window); the red dotted line indicates stimulus onset, and the whisker stimulus consisted of a 500 ms deflection train; *inset (green box)* indicates the analysis window used to quantify evoked responses. **f.** Baseline-corrected whisker-evoked multiunit activity (MUA) quantified from the analysis window indicated in **e** and compared between vehicle and CTZ epochs across stimulus amplitudes (pooled across low, medium, high trials). **g.** Trial-level regression relating late firing to early firing across recording channels in LMC and opsin–expressing mice. **h.** Same analysis as in g, performed in LMC-expressing mice lacking opsins.

We first tested whether luciferin-driven bioluminescence alone was sufficient to enhance spontaneous cortical activity in the absence of external light stimulation. By expressing CheRiff in vSI pyramidal neurons, we predicted increased MUA either through external LED stimulation or by luciferin-driven bioluminescence. CTZ application enhanced the MUA firing rates in opsin-expressing mice relative to the null distribution derived from mice lacking opsin expression (**Fig. 6c**, No opsin: 21.38 Hz vs opsin: 50.20 Hz two-sided, unpaired Wilcoxon rank-sum test, p= 3.159 ×10⁻¹¹). At the end of each recording session during CTZ perfusion, a brief sinusoidal LED stimulus (8 Hz, 500 ms) was delivered to confirm optogenetic responsiveness (**Fig. 6d**). LED-evoked MUA increased in opsin-expressing mice relative to the null distribution derived from the control cohort (No opsin: 2.456 Hz vs opsin: 86.420 Hz; two-sided, unpaired Wilcoxon rank-sum test, p= 7.6 ×10⁻¹⁷). Together, these results are consistent with bioluminescent optogenetic activation of CheRiff selectively enhancing spontaneous cortical activity *in vivo*.

Following our observation that LMC enhances cortical activity during luciferin perfusion, we next tested whether LMC can rapidly amplify sub-second neural responses to sensory inputs. Consistent with BLADe activation of CheRiff, we tested for relationships between enhanced neural activity and vibrissae stimulation by comparing MUA between vehicle and CTZ epochs. We delivered 10 vibrissae deflections at 20 Hz over a 500 ms stimulus window, with deflections randomized in amplitude. We observed evoked spikes within ∼10–20 ms in vSI following low-amplitude vibrissae deflections, consistent with known sensory latencies in vSI and validating our electrode placement (**Fig. 6e**). To isolate BLADe-dependent modulation of sensory-evoked cortical activity, we analyzed whisker-evoked multiunit activity (MUA) in a 500 ms post-stimulus window. We then computed the CTZ-induced change in stimulus-locked MUA (POST–PRE) for each channel and compared these responses between CheRiff-expressing and control mice. Opsin-expressing mice displayed enhanced evoked MUA activity in the presence of CTZ following continuous vibrissae deflections, consistent with BLADe dependent recruitment of sensory-evoked cortical responses (**Fig. 6f**, p= 2.65 ×10^−17^, two-sided, unpaired Wilcoxon rank-sum test). In control mice, baseline-corrected evoked responses were centered near zero and showed a slight bias toward decreased activity, indicating nonspecific CTZ-related modulation that was not contingent on opsin expression. Together, these findings indicate that LMC rapidly enhances stimulus-locked cortical activity by linking ongoing neural activity to additional optogenetic drive, leading to increased sensory evoked responses.

We next asked whether the activity-dependent gain observed in the patch clamp recordings with excitatory opsins extends to the circuit level *in vivo*. In CheRiff-expressing mice (**Fig. 6g**), luciferin perfusion produced a tonic increase in late activity (β_1_= 0.30, CI [0.28, 0.32], p= 1.29 ×10^−19^, β₂= 7.01, CI [5.16, 8.85], p= 9.998 ×10^−14^). After controlling for this optogenetic effect, luciferin perfusion allowed sensory-evoked responses to be amplified as a function of early spiking activity, such that trials with greater early activity exhibited larger responses during the late epoch (β₃= 0.0647, CI [0.044, 0.086], p= 1.721 ×10^−9^). This corresponded to a 22% enhancement of late-phase firing (β₃/β₁ = 0.0647/0.301). In contrast, mice not expressing CheRiff (**Fig. 6h**) showed no evidence of gain modulation (β₃ = −0.024, 95% CI [−0.061, 0.012], p = 0.189), corresponding to a negligible 4% decrease in slope. Luciferin perfusion produced a non–opsin-specific tonic suppression of late activity (β_1_= 0.6031, CI [0.568, 0.638], p= 8.38 ×10^−25^, β₂= −5.88, CI [−8.12, −3.64], p= 2.8 ×10⁻⁷). Application of CTZ revealed a multiplicative gain effect in late spiking activity as a function of early firing selectively in LMC and CheRiff co-expressing mice (**Supplementary Fig. S6**).

Collectively, these findings are consistent with LMC enabling cell-autonomous, activity-gated control of sensory-evoked cortical activity *in vivo*. Through co-expression of LMC with light sensitive ion channels, BLADe photoactivation offers a genetically encoded strategy for sensing and rapidly (sub-second) manipulating stimulus locked neural dynamics.

## Discussion

In this study we showed evidence in support of a bioluminescent activity-dependent (BLADe) platform that utilizes a luciferase split by a sensing moiety to drive a photoreceptor, allowing integration of intracellular biochemical fluctuations with cellular outcomes. For an example application, we engineered a Ca²⁺ dependent luciferase and applied it in two scenarios, the neuronal activity driven activation of a) a light sensing transcription factor and of b) a light sensing ion channel.

We split the *Gaussia* luciferase variant sbGLuc and inserted a Ca²⁺ sensing domain composed of the M13 peptide from YC3.6 and calmodulin from GCaMP6f to generate lumicampsin (LMC). When comparing LMC to other bioluminescent Ca²⁺ sensors we identified critical design constraints for enabling activity dependent control of photosensing molecules: at resting intracellular Ca²⁺ levels, the presence of luciferin should generate minimal light output – ideally below the activation threshold of photoreceptor chromophores. Furthermore, elevation of intracellular Ca²⁺ must produce a sufficient increase in bioluminescence to cross this threshold and initiate photoreceptor activation.

Several excellent bioluminescent Ca²⁺ sensors have been developed by splitting luciferase enzymes and inserting Ca²⁺ sensing moieties^16,18–20,28–30^, and new sensors for reporting various intracellular signals are being developed. Because bioluminescent probes have historically been far dimmer than fluorescent probes, most efforts have focused on maximizing light output. As a result, existing bioluminescent Ca²⁺ sensors successfully report Ca²⁺ fluctuations, but with Ca²⁺ dependent increases in signal riding on top of high baseline light emission. For many imaging applications, baseline bioluminescent signal in the absence of high Ca²⁺ is acceptable.

However, this is incompatible with the BLADe approach, as the background light emission by itself is already high enough to activate a photoreceptor. Thus, an excellent bioluminescent sensor cannot be automatically assumed to be a suitable driver for BLADe applications. To avoid unintended photoactivation, the split luciferase must produce very low levels of light emission at rest –when luciferin is present but intracellular calcium is low. Under these conditions, the luciferase enzyme should remain incompletely reconstituted and emit minimal light. Only upon calcium binding—or the presence of a specific co-factor needed for enzyme reconstitution—should bioluminescence steeply increase to a level sufficient for activating nearby photoreceptors. Comparison to other bioluminescent Ca²⁺ sensors revealed this critical difference in SNR for the LMC construct used here to promote the BLADe paradigm. Nonetheless, here we tested only a limited number of available sensors and there might be suitable sensors among the already available ones. Furthermore, available sensors with excellent performance in imaging applications should continue to be used for such but would require further analysis if they were to be considered as drivers of optogenetic actuators.

BLADe depends on conditional light emission that stays *low* at baseline and *rises steeply* when the target co-factor is present. LMC satisfied this criterion but is very dim and thus of limited broad use, which prompted the systematic development of a novel Ca²⁺ dependent luciferase with low baseline signal and high peak brightness, CaBLAM (Ca²⁺ BL Activity Monitor)^31^. Different from the *Gaussia* luciferase based LMC, CaBLAM was evolved from *Oplophorus gracilirostris* luciferase ^32–35^. Key differences to LMC in the CaBLAM sensor design are the choice of a split in the loop that joins the final two β-strands for insertion of the Ca²⁺-sensing domains, a different M13-CaM pair, an N-terminal fluorescent partner for FRET enhancement and restricting Ca²⁺ readout to the BL channel, and extensive affinity tuning that generated variants covering much of the physiological range of Ca²⁺ concentrations. While LMC served to demonstrate the BLADe concept and to reveal key aspects of BLADe, CaBLAM’s superb features position it as the Ca²⁺ indicator of choice for future iterations of BLADe, superseding LMC. Moreover, the molecular evolution of CaBLAM incorporates a new luciferase scaffold, SSLuc (Sensor Scaffold Luciferase)^31^ that is highly amenable to insertion of sensor domains, making it a flexible starting point for engineering an array of new BLADe indicators that couple other biochemical events to light output for tailored optogenetic output.

In previous studies, two approaches have been used for integrating intracellular event sensing with molecular outcomes by coupling bioluminescent light emission to downstream processes. In the first, interaction between an intact luciferase and a light sensing protein is designed to be contingent on an intracellular event, such as protein-protein interaction (SPARK2)^36^, Ca²⁺ increase (FLiCRE)^37^, or GPCR activation (LOVTurbo)^38^. The second approach uses a two-component split luciferase construct (LuCID)^39^, where two halves of a split luciferase were fused to two independent parts sensing Ca²⁺ (CaM and CaM binding protein). LuCID was also used to couple protein-protein interaction to transcription, requiring new constructs that fuse the two separate parts of the luciferase to two separate sequences detecting protein interaction^39^. The above systems require multiple components, which may be a feature of a tool in some applications but requires a lot of engineering for expanding its use to different contexts. In contrast, BLADe is a highly modular tool that uses a single molecule containing both halves of the luciferase split by a sensing moiety. Intracellular events that can be integrated with optogenetic target effects are only limited by which sensors can be designed. And the same bioluminescent sensor can be coupled with, in principle, any optogenetic actuator with matching excitation wavelength.

These previous approaches of bioluminescent sensing and modulating, i.e. integrating, have been applied to drive diverse cellular outcomes, including transcription of fluorescent reporters to label cells that experienced particular intracellular events, the transcription of actuators to allow manipulation of cells, and proximity labeling of nearby proteins. Bioluminescence has not been used previously for coupling intracellular Ca²⁺ to opsin activation for changing the same cell’s membrane potential. However, a Ca²⁺ dependent luciferase has been used for synaptic transmission of photons from an activated presynaptic neuron to a channelrhodopsin-expressing postsynaptic neuron (PhAST)^40^. This application of bioluminescence-mediated cell-to-cell communication supports the potential for using Ca²⁺ dependent bioluminescence emission for activating photoreceptors, even over some distance without tethering.

Bioluminescence has been used for coupling intracellular acidification to opsin activation for hyperpolarizing the cell^41^. Here, bioluminescence emitted from an intact luciferase was applied to couple a sensor for neuronal hyperactivity, i.e. acidification, with a molecular actuator capable of switching off neuronal activity in an all-molecular negative feedback loop at the single neuron level^41^. To generate a pH sensitive inhibitory luminopsin (pHIL), three proteins were fused: a hyperpolarizing opsin (eNpHR3,0), a pH sensing fluorescent protein (E2GFP), and a *Renilla* luciferase. Through a two-step resonant energy transfer cascade triggered by acidification in the presence of luciferin, pHIL translates the hyperactivity-induced intracellular acidosis into silencing of neuronal activity. Cell-autonomous hyperpolarization was demonstrated *in vivo* by mitigating acute tonic-clonic seizures induced by pilocarpine and by counteracting audiogenic seizures in the PRoline-Rich Transmembrane protein 2 knockout (PRRT2 KO) mouse, a model of genetic epilepsy. While pHIL is designed to specifically convert neuronal hyperactivity into silencing, the BLADe approach includes and transcends this application of real-time feedback. Co-expression of a Ca²⁺ BLADe integrator with an inhibitory or excitatory opsin results in neural activity-driven hyperpolarization or additional depolarization, respectively.

The general motivation for developing closed-loop systems for modifying cells based on their intracellular activity has led to a number of activity dependent regulatory approaches. The drive for developing such approaches has been most prolific in neuroscience, challenged by the question of how to identify and study the ensemble of active neurons underlying a specific behavior and to therapeutically control selected subsets of overactive neurons. Some of the current solutions are based on implanted hardware and computer-based detection algorithms to conduct detection and deliver control. For example, signals are being tracked using electrodes, and patterns detected by computer algorithms trigger countermanding stimulation through implanted electrodes or fiber optics for electrical or optogenetic regulation^42–44^. However, these methods require chronic implants to detect patterns and deliver stimuli, and electrical stimuli impact all adjacent cells. The desire to replace problematic “externally closed-loop” systems by “internal closed-loop sensor-actuators” working at the level of individual neurons motivated the development of tools for all-molecular feedback regulation. Several approaches were developed for coupling intracellular events to transcription of actuators using calcium sensitivity and engineered TEV protease^45–47^. Cal-Light^45^ and FLARE^46^ sense Ca²⁺ and are temporally gated by light exposure to drive transcription of actuators. Other systems, such as TRAP^48^ and CANE^49^, use immediate early gene expression as a proxy for neuronal activity to drive transcription. Refinements of these concepts have led to powerful preclinical tools, for example by using an immediate early gene promoter to drive the expression of an engineered potassium channel in hyperactive neurons for as long as they exhibit abnormal activity^50^. This activity-dependent gene therapy was effective for on-demand cell-autonomous treatment for epilepsy in mice. In contrast to sensor-actuator coupling via transcription, other approaches allow integration of biochemical events with cellular outcomes in real time. Again, in the context of epilepsy therapy, biochemical closed-loop treatments were achieved by employing pathologically elevated glutamate or acetylcholine to activate engineered neurotransmitter-gated chloride and glycine channels, respectively, to effectively control seizures in rodents^51,52^.

Conceptually, the BLADe platform encompasses these above applications by allowing sensing of fluctuations of any component for which a bioluminescent sensor can be designed and integrating these with a variety of optogenetic actuators for a wide spectrum of cellular outcomes. The BLADe design with a single-component sensor is very straightforward, different from several of the above systems that require more complicated molecular designs, increasing the potential for failure points in distinct cell types. Moreover, as the sensor can also be imaged, BLADe enables simultaneous monitoring of sensor-actuator interactions. The need for luciferin application places BLADe always under experimenter control. At the same time, it allows for tuned effects through variation in luciferin concentrations, its route of administration with different kinetics for intravenous versus intraperitoneal versus intraventricular application, use of specific luciferins with short-term flash versus long-term glow kinetics, and timing of luciferin administration throughout a behavioral paradigm.

Our study has important limitations. While our data are consistent with LMC integrating Ca²⁺ fluctuations to drive transcription and change membrane potential at the functional level, i.e. activity- and history-dependent coupling of cellular state change to output, we do not show a causal biochemical mapping between intracellular Ca²⁺ and cellular output. Such formal proof would require experiments employing Ca²⁺ chelators and Ca²⁺ titrations to establish a quantifiable causal relationship between Ca²⁺ levels and optogenetic output. However, this limitation does not negate the functional evidence from our *in vitro* and *in vivo* data that demonstrate BLADe coupling past activity to future output in real time. In fact, we show integration across two temporal regimes. First, in the opsin patch and *in vivo* sensory evoked data, the β₃ interaction demonstrates that prior activity changes how subsequent activity is generated within a single stimulus epoch—i.e., real-time, sub-second integration. Second, in the transcriptional experiments the readout reflects prior cellular activity, as shown in both our HeLa and mouse vSI experiments, where chemically enhancing intracellular activity (e.g., histamine or NMDA) leads to produce corresponding changes in gene expression. Nevertheless, the above biochemical studies should be done with the next generation of suitable Ca²⁺ sensors. Another set of experiments should be a titration of Ca²⁺ sensor expression as overexpression of luciferase-based sensors may lead to nonlinear signal behavior. While LMC served to establish the design requirements for Ca²⁺ sensors to serve as integrators and allowed to demonstrate applications of the BLADe approach, its light emission is very dim. For going forward, a new Ca²⁺ sensor suitable for BLADe applications superseding LMC has recently been developed^31^. In future work, the BLADe platform could be improved and extended in a number of ways. The luciferase can be split by different sensing domains whose light activation can be coupled to a wide range of light-sensing actuators, providing experimental control over diverse molecular outcomes depending on the user’s goal. Further developed and improved versions of these tools are expected to have broad applications. In basic research, this paradigm allows investigators to selectively detect, and then suppress or amplify, specific biological events, a unique approach to understanding the role of those processes. In potential therapeutic applications it allows sensing and discontinuing aberrant activity before it can cause harm, or stopping pathological processes from continuing, or amplifying and rescuing failing processes, without the use of implanted devices and regulated by simple peripheral injection of a luciferin.

## METHODS

### Chemicals

All chemicals, including Ionomycin and Histamine, were purchased from Sigma. The Calcium Calibration Buffer Kit #1 from Life Technologies was used to prepare Zero Ca²⁺ buffer and 39 µM Ca²⁺ buffer.

### Luciferins

For *in vitro* experiments, Coelenterazine (CTZ) (NanoLight Technology; Pinetop, AZ; NanoLight # 303) and Coelenterazine h (hCTZ) (NanoLight # 301) were dissolved in NanoFuel solvent (NanoLight # 399) and stored in 50 mM stock solutions at -80 °C. CTZ, hCTZ, or NanoFuel Solvent (for vehicle controls) were freshly diluted in culture medium for final concentrations. D-luciferin potassium salt was purchased from GoldBio (USA), dissolved in water and stored as 60 mg/ml stock aliquots at -80°C. For use in plate readings, stock solutions were diluted in OptiMEM, then further diluted into wells for 150 µg/ml final concentration. For *in vivo* experiments, water soluble Coelenterazine (Nanolight #3031) was freshly prepared by dissolving in sterile water to a stock concentration of 1.6 mM for lateral ventricle injection to yield a final concentration of 200 μM and to a stock concentration of 2.36 mM for dilution in ACSF to yield a final concentration of 20 μM.

### Plasmids

A pcDNA3.1 backbone with the CMV promoter was used to insert gBlocks (IDT) that encode for N-sbGluc, CaM-M13 variants, C-sbGluc and p2a dTomato by Gibson cloning (New England Biolabs HiFi DNA Master Mix). Coding sequences from plasmids for GCaMP6f, hGtACR2, ChR2(C128S), and CheRiff (Addgene) and for BlueCaMBI and GLICO (synthesized by Genscript) were cloned into the pcDNA3.1-CMV backbone. Expression plasmids for VP-EL222 and 5xC120-FireflyLuc were kindly provided by Dr. Kevin Gardner, CUNY, New York, NY, and for GeNL(Ca2+) by Dr. Takeharu Nagai (Addgene). For viral vectors, coding sequences were cloned into a pAAV vector downstream of a CAG or hSyn promoter, or into a pAAV-EfIa-DIO construct. For list of plasmids used see Supplementary Table 1.

pcDNA3.1-CAG-VP-EL222 and pcDNA3.1-5xC120-FireflyLuc were kindly provided by Dr. Kevin Gardner, CUNY, New York, NY pAAV.Syn.GCaMP6f.WPRE.SV40 was a gift from Douglas Kim & GENIE Project (Addgene plasmid # 100837 ; http://n2t.net/addgene:100837 ; RRID:Addgene_100837) GeNL(Ca²⁺)_520/pcDNA3 was a gift from Takeharu Nagai (Addgene plasmid # 85204 ; http://n2t.net/addgene:85204 ; RRID:Addgene_85204) pLenti-CaMKIIa-hChR2(C128S)-EYFP-WPRE was a gift from Karl Deisseroth (Addgene plasmid # 20294 ; http://n2t.net/addgene:20294 ; RRID:Addgene_20294) pFUGW-hGtACR2-EYFP was a gift from John Spudich (Addgene plasmid # 67877 ; http://n2t.net/addgene:67877 ; RRID:Addgene_67877) pAAV-hSyn-CheRiff-eGFP was a gift from Adam Cohen (Addgene plasmid # 51697 ; http://n2t.net/addgene:51697 ; RRID:Addgene_51697)

### AAV

AAV2/9 preparations were generated by triple lipofection of HEK293-FT cells and harvesting viral particles as previously described^5^. Briefly, triple plasmid transfection in HEK293 cells (Agilent, Cat # 240073-41) was performed. The helper plasmid pAd delta F6 (24 μg), respective pAAV-hSyn plasmids (12 μg), and the serotype plasmid AAV2/9 (20 μg) were transfected, using Lipofectamine 2000, into subconfluent HEK293 cells grown in 10 cm culture dishes. Virus was purified from supernatant and cells with an aqueous two-phase system after 72 hrs, dialyzed at 4°C overnight against PBS (Ca2+ and Mg2+ free). The next morning, virus was concentrated in Amicon Ultra-0.5 mL Centrifugal Filters. Q-PCR targeting the WPRE element was used to determine viral titers.

### Animals

All mouse experiments described in this study were approved by the Central Michigan University Institutional Animal Care and Use Committee (IACUC) and conducted in accordance with all relevant guidelines and regulations, including the US National Research Council’s Guide for the Care and Use of Laboratory Animals, the US Public Health Service’s Policy on Humane Care and Use of Laboratory Animals, the Guide for the Care and Use of Laboratory Animals, and the ARRIVE guidelines (Animal Research: Reporting of In Vivo Experiments, PLoS Bio 8(6), e1000412,2010) for reporting animal research. Mice were group-housed in ventilated cages under 12-hour reverse light cycle, provided with tap water and standard chow and allowed to feed ad libitum. C57BL/6 (JAX #000664), Emx1-Cre (JAX# 005628), and Swiss Webster (Charles River) mice of both sexes were used. Mice were euthanized by CO_2_ asphyxiation followed by cervical dislocation in accordance with the above guidelines for animal euthanasia.

### Cell Culture and Cell Transformation

To avoid any ambient light contamination, all tissue culture work was carried out in a dark room with a red light source; this room also housed the cell culture incubator. Human embryonic kidney fibroblasts (HEK293, RRID:CVCL_0045) and human cervical adenocarcinoma cells (HeLa, RRID:CVCL_0030) were purchased from the American Type Culture Collection (HEK293: ATCC, CRL-1573, Lot #70039815; HeLa: ATCC, CCL-2, Lot #70037076), expanded, and aliquots at low passage numbers were frozen. Cells were grown in Gibco DMEM media supplemented with 10% Fetal Bovine Serum, 1% Glutamax, 0.5 % Pen-Strep, 1% Non-Essential Amino Acids and 1% sodium pyruvate. Cells were cultured at 37°C and 5% atmospheric carbon dioxide. Cells were transfected with Lipofectamine 2000 or 3000 in accordance with the manufacturer’s instructions.

Primary E18 rat cortical and hippocampal neurons were prepared from tissue shipped from BrainBits (Transnetyx) following the vendor’s protocol. Neurons were grown in Gibco Neurobasal Media supplemented with 2% B27 supplement and 0.1% gentamycin and 1% Glutamax. Neurons were nucleofected with plasmids of interest using the Lonza 2b Nucleofector and Rat Neuron Nucleofector Kit (Lonza, VPG-1003) according to the manufacturer’s instruction, then seeded. Or neurons were nucleofected, seeded, then transduced with AAV the next day. Neurons were seeded on PDL-coated glass coverslip (Neuvitro) in 12-well tissue culture plates (1 × 10^5^ neurons per well) or were plated on the electrode area of 1-well MEA dishes (60MEA200/30iR-Ti; Multi Channel Systems, Germany) coated with PEI (0.1%) and laminin (50 µg/ml) (1×10^5^ neurons/10 µL/well) as described in detail in Prakash 2020^26^.

Primary cardiomyocytes were harvested from E18 Swiss Webster mice, papain digested and plated on 35mm glass bottom dishes (MatTek) for real time bioluminescence imaging. Primary cardiomyocytes were grown in Claycomb Media supplemented with 10% HL-1 screened FBS, 100 µg/mL penicillin/streptomycin, 0.1mM norepinephrine and 2% Glutamax. Cells were nucleofected with plasmids of interest using the Lonza 2b Nucleofector and Rat cardiomyocyte Nucleofector Kit (Lonza, VAPE-1002) according to the manufacturer’s instruction.

### Microscopy

Initially, all transfected cells were imaged using a Zeiss Axio Observer A1 microscope with a LD A-Plan 20x/0 air objective and a Hamamatsu Orca Flash 4.0 CMOS camera to confirm expression by fluorescence microscopy at 50 ms exposure. Imaging of brain sections were done on an Olympus Fluoview 300 CLSM confocal microscope with a 60x/1.3 Oil objective.

### Bioluminescence Imaging

For photon counting HEK293 and HeLa cells were seeded on Costar 96 well clear bottom white plates and luminescence was measured in a SpectraMax® L luminometer (Molecular DevicesTM). For real-time bioluminescence imaging we used a Zeiss Axio Observer A1 microscope with a Fluar 40x/1.3 Oil objective with either a Hamamatsu Orca Flash 4.0 CMOS camera or an Andor iXon Ultra 888 EM-CCD camera with their respective softwares. HEK293 and HeLa cells were imaged in imaging solution (CaCl2 1.25 mM, HEPES 19.7 mM, KCl 4.7 mM, KH2PO4 1.2 mM, MgSO4 1 mM, NaCl 130 mM, dextrose 0.5, pH 7.2-7.4) with either 100 µM or 12.5 µM CTZ. Primary cardiomyocytes were imaged in Tyrode’s solution (NaCl 137 mM, KCl 2.7 mM, MgCl2 1 mM, CaCl2 1.8 mM, Na2HPO4 0.2 mM, NaHCO3 12 mM, D-glucose 5.5 mM, pH 7.2-7.4) supplemented with 10 µM norepinephrine and 100 µM CTZ. Primary E18 rat hippocampal neurons were imaged in artificial cerebrospinal fluid, containing: 121 mM NaCl, 1.25 mM NaH2PO4, 26 mM NaHCO3, 2.8 mM KCl, 15 mM d(+)-glucose, 2 mM CaCl2, 2 mM MgCl2 (maintained at ∼37 °C, pH 7.3-7.4, continuously bubbled with 95% O2/5% CO2 vol/vol). All imaging was conducted in the RC-49MFSH heated perfusion chamber where the temperature was continuously monitored (Warner Instruments).

### Electrophysiology recordings from primary neurons

#### Whole Cell Recordings

Neurons (rat E18, cortical) were co-nucleofected with LMC4f and either an inhibitory opsin, hGtACR2, an excitatory opsin, ChR2(C128S), or a “Dud” opsin, ChR2(C128S)-E97R-D253A. Post nucleofection, neurons were plated on laminin-coated glass coverslips (15mm, Neuvitro) in culture medium consisting of Neurobasal Medium (Gibco # 21103-049), B-27 supplement (Gibco # 17504-044), 2 mM Glutamax (Gibco # 35050-061), and 5% Fetal Calf Serum (FCS). The following day, the medium was replaced with serum-free medium (NB-Plain medium). Half of the medium was replaced with fresh NB-Plain medium every 3–4 days thereafter. Neurons were used for whole cell patch clamp recordings between DIVs 21–25. For patch clamp recording, a coverslip was transferred to a recording chamber mounted on an upright microscope (BX51WI, Olympus) and perfused with aCSF containing (in mM): 121 NaCl, 2.8 KCl, 1 NaH_2_PO_4_, 26 NaHCO_3_, 2 CaCl_2_, 2 MgCl_2_ and 15 D-glucose (310 mOsm/kg, pH 7.3-7.4) at a rate of 1.5 ml/min. All solutions were bubbled with a gas mixture of 95% O_2_ and 5% CO_2_. Whole-cell patch clamp recordings were performed using a Multiclamp 700b amplifier and Digidata 1440 digitizer together with the pClamp recording software (Molecular Devices). Borosilicate glass micropipettes were manufactured using a PC-100 puller (Narishige) and had resistances of 3–5 MΩ. In current clamp recordings, pipettes were filled with intracellular solution containing (in mM): 130 K-gluconate, 10 KCl, 15 HEPES, 5 Na_2_-phosphocreatine, 4 Mg-ATP and 0.3 Na-GTP (310 mOsm/kg, pH 7.3). The aCSF was supplemented with D-AP5 (50 µM), CNQX (15 µM) and picrotoxin (100 µM) to block fast glutamatergic and GABAergic synaptic transmission. Rs was compensated by using bridge balance. Firing frequencies were calculated from the total number of action potentials produced per unit time during depolarizing current injections (10 sweeps, 1.5 sec duration) in episodic stimulation acquisition mode, at a membrane potential of -70mV, before and after CTZ (100 µM) or vehicle treatments.

To quantify activity-dependent changes in spike generation during sustained depolarization, we computed a spike count fractional difference for each trial. For each 1-s depolarizing current injection, spikes were binned into the first and second halves of the stimulus (0–0.5 s and 0.5–1.0 s). The number of spikes in the second half was subtracted from the number of spikes in the first half and normalized by the total spike count for that trial. This metric ranges from +1, indicating all spikes occurred in the first half of the stimulus, to −1, indicating all spikes occurred in the second half, with 0 reflecting an even distribution across the stimulus. This analysis was applied on a per-trial basis before and after CTZ or vehicle treatment to assess how LMC-dependent opsin activation altered the temporal distribution of action potentials during depolarization.

#### MEA Recordings

Once the neurons (rat E18, cortical) were matured (DIV14-DIV19), recordings were conducted using a MEA2100-Lite-System (Multichannel Systems, Germany). As described in detail in Prakash et al.^26^, raw extracellular signals were acquired via the MCS system and stored in HDF5 format. All-trans retinal (R2500; Sigma- Aldrich, St. Louis, MO) was added to the culture medium to 1 μM final concentration before electrophysiological recordings. Prior to recording, all reagents were pre-warmed to 37 °C. A micropipette was used to add reagents with the reagent drop gently touching the liquid surface, creating a time-locked artifact in the recordings. Recordings were carried out with a sample rate of 10 kHz. Raw samples were read directly from the HDF5 files and analyzed in native ADC units. Conversion to physical units was defined by MCS channel metadata (ADZero, ConversionFactor, Exponent) according to the manufacturer’s specification; for the recordings analyzed here, this corresponded to a scaling of 0.0596 µV per ADC count.

For visualization and spike detection, voltage traces were temporally aligned to the timestamp of chemical application (CTZ or vehicle). This timestamp was identified by a large, stereotyped amplitude artifact produced by direct addition of solution to the MEA which provided a within well experiment-wide temporal reference. A symmetric time window (±1000 ms) centered on this chemical event was extracted, defining pre- and post-application analysis periods. Chemically aligned voltage traces were high-pass filtered to isolate spiking activity from slower components of the extracellular signal (e.g., baseline drift and low-frequency fluctuations). Filtering was performed using a 4th-order Butterworth high-pass filter with a 300 Hz cutoff, applied with zero-phase forward–reverse filtering (filtfilt) to preserve spike timing.

Detection thresholds were defined as five times the baseline noise level, where baseline noise was estimated from the pre-chemical window using the median absolute deviation (MAD).In brief, noise was computed as 1.4826 × median(|yb − median(yb)|), where yb denotes the filtered baseline signal for each channel, and the detection threshold was set to 5.0 × noise. Negative spikes were identified as threshold-crossing deflections in the filtered extracellular signal using peak detection. Detected events were constrained by a refractory period of 1.0 ms and a minimum spike width of 0.3 ms; events occurring within 0.3 ms of one another were merged to prevent double counting. Only spikes occurring within the post-chemical window were retained for downstream analyses and were treated as multi-unit activity (MUA). Similar to *in vivo* recordings, MUA was estimated by converting raw spike times into instantaneous firing rates using 1 ms bins.

#### Transcription *in vitro*

HeLa cells were used to test whether LMC-driven light emission could activate transcription through the light-responsive transcription factor EL222. Cells were plated in 6-well plates and transfected at ∼80–90% confluency using Lipofectamine 2000 (Thermo Fisher). Each well received a total of 2 μg DNA, composed of the following: 666 ng EL222 (light-sensitive transcription factor), 333 ng 5×C120-FLuc (EL222-responsive firefly luciferase reporter), and 1000 ng LMC4f-dTomato (calcium-dependent luciferase). Parallel wells were transfected with other calcium-dependent bioluminescent sensors (GeNL, GLICO, BlueCaMBI) and matched FLuc reporters.

Three hours after transfection, cells were trypsinized (300 μL trypsin per well) and replated into PDL-coated, white-walled, clear-bottom 96-well plates (∼100,000 cells per well). Cells were allowed to adhere for 9–12 hours before stimulation. For transcriptional activation, 100 μL of stimulation buffer was added per well (1:1 with cell media) for a final well volume of 200 μL. The stimulation solution contained 100 μM CTZ or hCTZ (depending on the sensor), with or without 10 μM histamine. Vehicle controls received no treatment. After stimulation, cells were incubated for 7–9 hours before transcriptional reporter readout.

For FLuc transcriptional measurement, activity was recorded 18 hours after stimulation using a luminometer. Prior to measurement, cells were washed and incubated in phenol red–free media (100 μL per well). D-luciferin (final concentration 150 μg/mL) was prepared from a frozen 60 mg/mL stock by serial dilution into Opti-MEM. For each well, 25 μL of 750 μg/mL D-luciferin working stock was injected into 100 μL of media. FLuc bioluminescence was recorded immediately after injection.

#### Transcription *in vivo*

To test bioluminescent dependent transcription *in vivo*, we co-injected three AAV constructs into the prefrontal cortex of anesthetized mice (isoflurane at a rate maintained between 2 – 3% and oxygen at 55 mL/min; SomnoSuite Low-Flow Anesthesia System, Kent Scientific Corporation): AAV9-hSyn-LMC4f-P2A-dTomato (Ca²⁺-sensitive luciferase), AAV9-CAG-EL222 (light-activated transcription factor), and AAV9-5×C120-EYFP (EL222-responsive reporter). Viral solutions were mixed prior to injection to a total volume of 1 μL per site, yielding final per-virus titers of: LMC4f (1 × 10^10^ GC), EL222 (0.7 × 10^10^ GC), and C120-EYFP (0.3 × 10^10^ GC). The injection was targeted to the left medial prefrontal cortex at +0.37 mm anterior to bregma, 3.0 mm lateral. Following injection, the craniotomy was sealed to minimize ambient light exposure during recovery.

At the time mice were expressing light-sensing constructs, experiments were conducted in a dark room illuminated with red light. After 7 days, mice received intraventricular (ICV) injections of one of three treatments: (1) NMDA + vehicle, (2) luciferin (CTZ) alone, or (3) luciferin + NMDA. These conditions were used to assess background transcription, spontaneous activity-driven transcription, and full BLADe-dependent transcriptional gating, respectively. Injections were performed using a 26-gauge needle and 10 μL Hamilton syringe, targeting the right lateral ventricle (AP: –0.5 mm, ML: –1.1 mm, DV: –2.0 mm from bregma). The luciferin was prepared fresh by dissolving coelenterazine (NanoLight #3031) in sterile water to a stock concentration of 1.6 mM, then when injected was assumed to be diluted 1:8 in CSF within the ventricle to yield a final injection concentration of 200 μM. NMDA was prepared at a working concentration of 75 ng/μL in sterile water. For combined delivery, 1 μL of NMDA solution (75 ng) was mixed with 4 μL of luciferin solution per injection (final 5 μL per mouse, injected over 5 mins). Solutions were prepared under red light and protected from exposure throughout. After injection, mice were removed from the stereotaxic apparatus and allowed to recover. Brains were collected for histological analysis ∼20 hours after ICV injection. Mice were anesthetized with isoflurane and transcardially perfused with PBS followed by 4% paraformaldehyde. Coronal brain sections were collected and imaged by confocal microscopy to quantify EYFP reporter expression. Confocal imaging of EL222 was performed using a rabbit anti-VP16 tag primary antibody (Abcam, ab4808), followed by a far-red Alexa Fluor 647-conjugated goat anti-rabbit secondary antibody (Invitrogen, A-21245) to minimize spectral overlap with dTomato fluorescence from LMC.

To analyze calcium- and light-dependent transcriptional activation *in vivo*, we quantified EYFP reporter expression in dTomato-labeled neurons across experimental groups, with the individual scoring sections blinded to experimental groups. Confocal images from all treatment conditions (NMDA-only, luciferin-only, luciferin + NMDA) were analyzed. First, dTomato+ images were pooled and segmented from maximum intensity projections using the MATLAB ‘cell-segm’ thresholding and size-filtering algorithm^53^, ensuring unbiased detection across all fields of view. The resulting binary masks were then applied to the corresponding EYFP images to extract raw per-cell reporter fluorescence from each dTomato+ neuron. Finally, cells were grouped by treatment, and the ratio of EYFP to dTomato fluorescence was computed per cell to normalize for expression variability.

#### Electrophysiology *in vivo*

To express LMC and CheRiff in excitatory cortical neurons, we injected AAVs into the left vibrissa primary somatosensory cortex (vSI) of Emx1-Cre mice (3–4 month old). AAV1-EF1α-DIO-LMC4f and AAV1-EF1α-DIO-CheRiff-EYFP were mixed 1:1 and delivered in a single injection (1 μL total) using a 33-gauge Nanofil syringe. Using a dental drill, a burr hole was made for AAV injection. The injection site was located 1.5 mm posterior and 3.0 mm lateral to bregma, and the needle was lowered to a depth of 500–1000 μm below the pial surface. Virus was infused at a rate of 50 nL/min over ∼20 minutes. Following injection, the needle was held in place for 10 min, then raised to 200 μm and held for an additional 10 min before full withdrawal to allow the virus to spread across the layers of the cortex. During the same surgery, a custom metal headpost was implanted for head fixation. After exposing the skull, a thin layer of clear Metabond was applied to stabilize the surface while keeping bregma and surface vasculature visible. The headpost was then affixed using additional Metabond, positioned off-center to leave vSI accessible. The short arm was angled over the left ear to allow right-side whisker stimulation. After injection and headpost implantation, the craniotomy was covered with mineral oil and sealed with Metabond. Animals recovered for at least 3 weeks before recordings.

After 2–3 weeks of recovery, mice were anesthetized with isoflurane (1.2–1.4%) and head-fixed. Again, these experiments were conducted in a dark room illuminated with red light. The craniotomy used for AAV injection was reopened and expanded to 1 mm to expose the underlying cortex and target the same vSI region as for the viral delivery. Throughout the procedure, the brain remained covered in carbogenated ACSF (95% O₂ / 5% CO₂) under constant perfusion. Animals were placed in a light-tight, electrically shielded chamber and maintained under anesthesia. Mice were secured in place to allow perfusion tubing to be positioned next to the craniotomy, allowing for constant ACSF inflow (for bioluminescence CTZ was diluted to 20 µM in ACSF). All macrovibrissae on the right side of the mouse were secured ∼3mm from the mystacial pad. A custom-made clamp was glued to a piezoelectric bender (Noliac CMBP09), positioned to move all macrovibrissae in the caudorostral direction, with a half-sine wave velocity profile with a rising phase (6ms) and a slower relaxation phase (20ms). On each trial, 20 Hz vibratory vibrissal stimulus trains (10 deflections, 500 ms) were delivered for 1000 trials total. On a trial by trial and randomized basis, the stimulus amplitude varied between 0 to maximal amplitude (∼1mm deflection). For each recording session, ACSF was constantly perfused for the first 500 trials and luciferin (CTZ) was then perfused for the remaining 500 trials.

All recordings in barrel cortex were conducted under light isoflurane anesthesia. Laminar probes consisted of single shank,32-channel silicon probes with a fiber optic 50 μm above the highest recording site (A1×32 Poly2-5mm-50s-177-OA32LP, Neuronexus Technologies; 0.15mm silver wire reference). Data was sampled at 30 kHz and passed through a digital amplifier (Cereplex-μ, Blackrock Microsystems), and directed through HDMI to the Cereplex Direct data acquisition box (Blackrock Microsystems). The 32-channel linear silicon probe was inserted perpendicularly into cortex using a micromanipulator at ∼10 µm/s until the top contact of the electrode disappeared beneath the pial surface. The probe was allowed to settle for ∼30–60 minutes before recording. Luciferin (coelenterazine; NanoLight #3031) was freshly prepared in sterile water (1 µg/mL) and diluted into ACSF to a final concentration of 20 μM. Following an initial recording period under vehicle ACSF, the perfusion line was switched to CTZ-containing ACSF to enable continuous bath application during the second half of the session. Flow rate and volume were monitored to maintain complete coverage of the craniotomy and probe without overflow.

Raw electrophysiological data were downsampled to 10 kHz. For each recording, electrode contacts with RMS values exceeding three times the interquartile range above the 75th percentile or below the 25th percentile across the 32-channel array were flagged as noise and removed for further analysis. All remaining electrodes were re-referenced to the common average^54^. MUA was estimated by converting raw spike times into instantaneous firing rates using 1 ms bins. For luciferin experiments, stimulus-aligned firing rates were extracted from trials before and after the defined luciferin application time.

For sensory stimulation experiments, MUA peristimulus time histograms (PSTHs) were constructed by aligning spike data to stimulus onset and binning at 1 ms. PSTHs were smoothed using a Gaussian kernel (window = 20 ms, SD = 3 ms) and averaged across trials. PSTHs were computed separately for pre- and post-luciferin trial blocks. Optogenetic responses to LED stimulation were quantified using a pre-stimulus baseline window (−500 to 0 ms) and a post-stimulus response window (0 to 500 ms). For each channel, the change in firing rate was defined as the difference between mean firing during the response window and baseline. A null distribution was constructed from no-opsin control recordings and used to define response thresholds. CTZ-evoked responses were quantified using the same baseline window by computing the change in firing rate between post-CTZ and pre-CTZ periods for each channel. A corresponding null distribution was generated from no-opsin control recordings using the same procedure and used for channel classification.

### Quantification of BLADe-dependent modulation of sensory-evoked activity

To isolate BLADe-dependent modulation of sensory-evoked cortical activity, whisker-evoked multiunit activity (MUA) was quantified on a trial-by-trial basis using a baseline-corrected metric. For each recording channel, peri-stimulus time histograms (PSTHs) were generated by time-locking spike times to stimulus onset within a 1500 ms window, pooling across stimulus amplitudes, and converting spike counts to firing rate (Hz) by scaling by the bin width. The alignment set stimulus onset to 0 ms across PSTHs and was used for visualization.

For each recording channel, spike trains were first aggregated across trials to compute firing rates aligned to stimulus onset. The mean firing rate during a 500 ms post-stimulus analysis window (500–1000 ms after stimulus onset) was then computed and baseline-corrected by subtracting the mean firing rate during the 500 ms immediately preceding stimulus onset (−500 to 0 ms) from the same channel. This channel-wise baseline subtraction normalizes evoked responses relative to ongoing activity within each recording channel, removing contributions from spontaneous firing, slow temporal drift, and global CTZ-induced changes in baseline excitability that are not time-locked to the sensory stimulus.

For each recording channel, CTZ-dependent modulation of sensory-evoked responses was quantified by computing the difference between baseline-corrected evoked firing rates measured during CTZ trials and those measured during vehicle trials. This within-recording subtraction isolates changes in stimulus-evoked activity attributable to CTZ within each channel after normalization to baseline firing. Therefore, the resulting channel-wise CTZ–vehicle difference served as the primary quantitative measure used to compare CTZ-dependent modulation of sensory-evoked activity between opsin-expressing and non-opsin control mice.

### Analysis of feed-forward effects in BLADe

For the linear regression, late evoked activity was modeled as a linear function of early evoked spiking activity and luciferin condition according to:

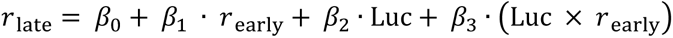

Here, (*r*_late_) is the late evoked firing rate measured in the late window (>200 ms after stimulation onset) while (*r*_early_) is the early evoked firing rate measured in the 0–200 ms window following stimulation onset. Luc indicates the absence (Luc = 0; vehicle) or presence (Luc = 1; luciferin, CTZ). β_0_ and β_1_ define the intercept and slope relating early to late activity in the absence of luciferin, whereas β_2_ captures a luciferin-dependent shift in the intercept (offset) and β3 captures a luciferin-dependent change in the slope relating early and late activity (gain). Early and late responses were pooled across trials, with regression performed at the level of individual neurons for patch-clamp recordings and at the level of individual recording channels for *in vivo* recordings. In this model, β₂ reflects a tonic, activity-independent shift in firing during the late phase window. As the model includes a tonic term (β₂), the interaction term (β₃) captures changes in the slope of the early–late relationship after accounting for activity-independent shifts. Thus, β₃ represents a luciferin-dependent change in the relationship between early and late spiking (gain). A positive β₂ indicates a tonic increase in late firing independent of early activity, whereas a negative value indicates tonic suppression. For β₃, positive values reflect multiplicative gain by strengthening the early–late relationship, whereas negative values reflect divisive neuromodulation.

### Statistics

Preprocessing and analysis of the data was performed with custom-written Matlab (R2020a, Mathworks) and Python scripts.

In **Fig. 1m**, maximal LMC luminescence responses (Max ΔRLU/RLU₀) with CTZ alone and glutamate+CTZ were compared using paired Wilcoxon signed-rank tests within the same neurons (same neuron, CTZ vs CTZ+glutamate). In **Fig. 2c**, histamine-specific changes in bioluminescence normalized to the luciferin baseline were compared across calcium sensors using a one-way ANOVA followed by Tukey’s post hoc multiple-comparisons test as an initial exploratory characterization of calcium-dependent luminescence responses. In **Fig. 3c**, overall differences across sensors were assessed using a Kruskal–Wallis test (non-parametric ANOVA), followed by targeted Wilcoxon rank-sum tests for hypothesis-driven comparisons involving LMC. In **Fig. 3g**, EYFP reporter/dTomato expression ratios from three independent conditions were compared using Wilcoxon rank-sum tests applied to all pairwise group comparisons, with Bonferroni correction for multiple testing (CTZ+NMDA vs CTZ vs NMDA). In **Fig. S3 and S4,** cumulative distributions of spike count fractional differences between the first and second halves of depolarizing current injections were compared using two-sample Kolmogorov–Smirnov tests across vehicle and CTZ perfusion conditions. In **Fig. S5,** luciferin-dependent changes in multiunit activity (MUA) were compared between inhibitory-opsin recordings with and without LMC co-expression using an unpaired Wilcoxon rank-sum test. In **Fig. 4b–d(iii)**, luciferin and vehicle action potential counts were compared using paired Wilcoxon signed-rank tests, with measurements paired within the same neurons across depolarizing steps under vehicle and CTZ conditions (same neuron, vehicle vs CTZ OR vehicle vs vehicle). In **Fig. 4b–d(iv)**, early and late spike counts were related using linear regression performed within individual neurons under vehicle and CTZ conditions. In **Fig. 5**, CTZ and vehicle responses were compared using paired Wilcoxon signed-rank tests, with each data pair corresponding to measurements from the same electrode recorded under vehicle and CTZ conditions at different time points on the same day (same electrodes, vehicle vs CTZ). In **Fig. 6 c,d, and f**, evoked firing changes in opsin-expressing and no-opsin channels were compared using an unpaired Wilcoxon rank-sum test. In **Fig. 6 g and h,** early and late evoked activity were related using linear regression performed at the level of individual recording channels under vehicle and luciferin conditions.

## Acknowledgments

We would like to thank all members of Bioluminescence Hub laboratories for their feedback, discussions, and thoughtful comments throughout the progression of this work. (http://www.bioluminescencehub.org/)

## Funding

This work was supported by National Institutes of Health grants R21MH101525, R21EY026427, U01NS099709, F99NS129170, National Science Foundation grants CBET-1464686, DBI-1707352, and the W.M. Keck Foundation.

## Author contributions

Conceptualization, AP, ELC, NCS, CIM, UH; Validation, AP, ELC, CIM, DL, NCS, UH; Formal Analysis, AP, ELC; Investigation, AP, ELC, MP, ADS, ZZ, MG-R, MOT; Data Curation, AP, ELC; Writing—Original Draft, AP, ELC; Writing—Review & Editing, ELC, NCS, CIM, UH; Visualization, AP, ELC; Supervision, CIM, NCS, UH; Project Administration, UH; Funding Acquisition, CIM, DL, NCS, UH.

## Data availability

The source data supporting the conclusions of this research is provided in this article and in the supplement. All datasets and custom scripts generated during these studies are freely available via the Brown Digital Repository (DOI pending).

## Competing interests

The authors have no conflicts of interest to declare.

## Notes

### Competing Interest Statement

The authors have declared no competing interest.

### Summary of Updates

Revision of main text (Results, Discussion, Methods), including new analyses of feed-forward effects in BLADe. Revision of main figures 1, 3, 4, 6. Revision of supplementary figures 1, 2, 3, 5, 6.

