## Supplementary Material for "A Bioluminescent Activity Dependent Platform, BLADe, for Converting Intracellular Activity to Photoreceptor Activation"

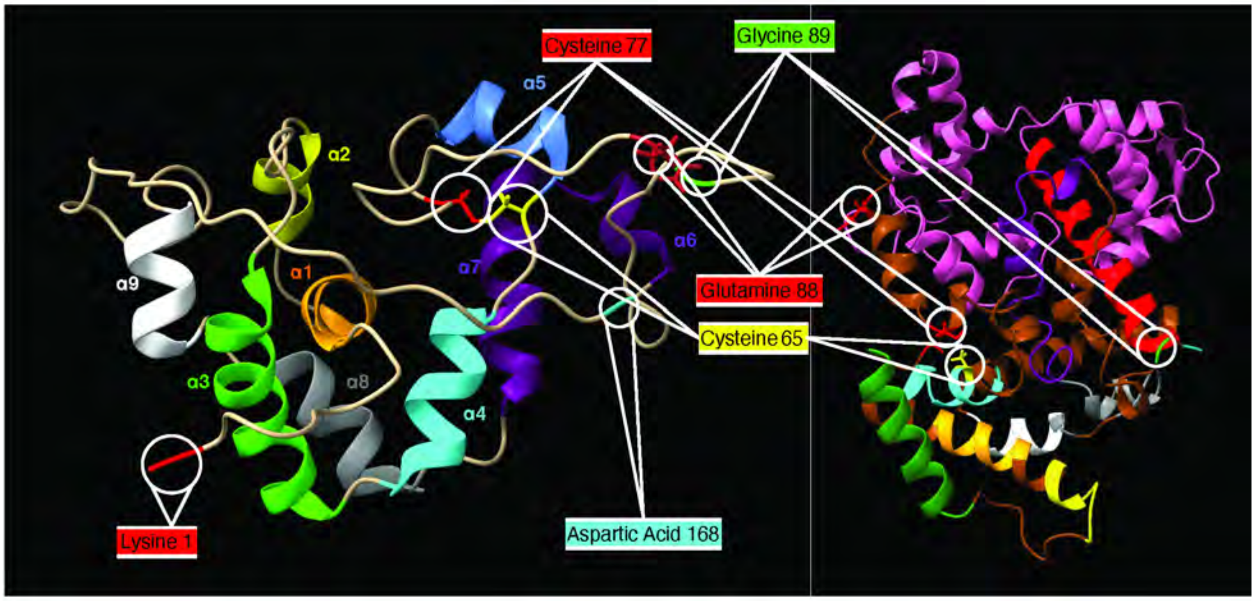


**Supplementary Figure S1. Ribbon diagrams of GLuc indicating 9 alpha helices**. Left: Of the 9 alpha helices α4 and α7 form the most rigid block of the luciferase stabilized by a disulphide bond between Cysteine 65 and 77. Right: The Calcium sensing moiety, CaM-M13 was inserted between Q88 and G89. In its unbound state, the CaM-M13 destabilizes the C65-C77 disulphide bond just by pushing the two halves away from each other. When bound to $Ca^{2+}$, the CaM-M13 pulls on both halves and stabilizes the C65-C77 disulphide bond to render the luciferase functional.

**a**

sbGluc(M43L_M110L):

K[1]PTENNEDFNIVAVASNFATTDLDADRGKLPGKKLPLEVLKELEANARKAGCTRGCLIC
LSHIKC[65]TPKMKKFIPGRC[77]HTYEGDKESAQ[88]G[89]GIGEAIVDIPEIPGFKDLEPLEQF
IAQVDLCVDCTTGCLKGLANVQCSDLLKKWLPQRCATFASKIQGQVDKIKGAGGD[168]

**b**

LMC: GLuc Signal Sequence; sbGluc N half; Linker #1; CaM;
Linker #2; M13; Linker #3; sbGluc C half; Alpha Helix

MGVKVLFALICIAVAEAK[1]PTENNEDFNIVAVASNFATTDLDADRGKLPGKKLPLEVLKELE
ANARKAGCTRGCLICLSHIKC[65]TPKMKKFIPGRC[77]HTYEGDKESAQ[88]GGGGTMADQLTEE
QIAEFKEEFSLFDKDGDGTITTKELGTVMRSLGQNPTEAELQDMINEVDADGDGTIDFPEF
LTMMARKMKYRDTEEEIREAFGVFDKDGNGYISAAELRHVMTNLGEKLTDEEVDEMIREAD
IDGDGQVNYEEFVQMMTAKGGKRRWKKNFIAVSAANRFKKISSSGALGSGGGG[89]GIGEAIV
DIPEIPGFKDLEPLEQFIAQVDLCVDCTTGCLKGLANVQCSDLLKKWLPQRCATFASKIQG
QVDKIKGAGGD[168]

**Supplementary Figure S2. Amino acid sequences of sbGLuc and LMC. a.** Amino acid sequence of sbGLuc with landmark mutations of sbGLuc versus wildtype GLuc color coded. **b.** Amino acid sequence of LMC with landmarks indicated by colors and underlines.

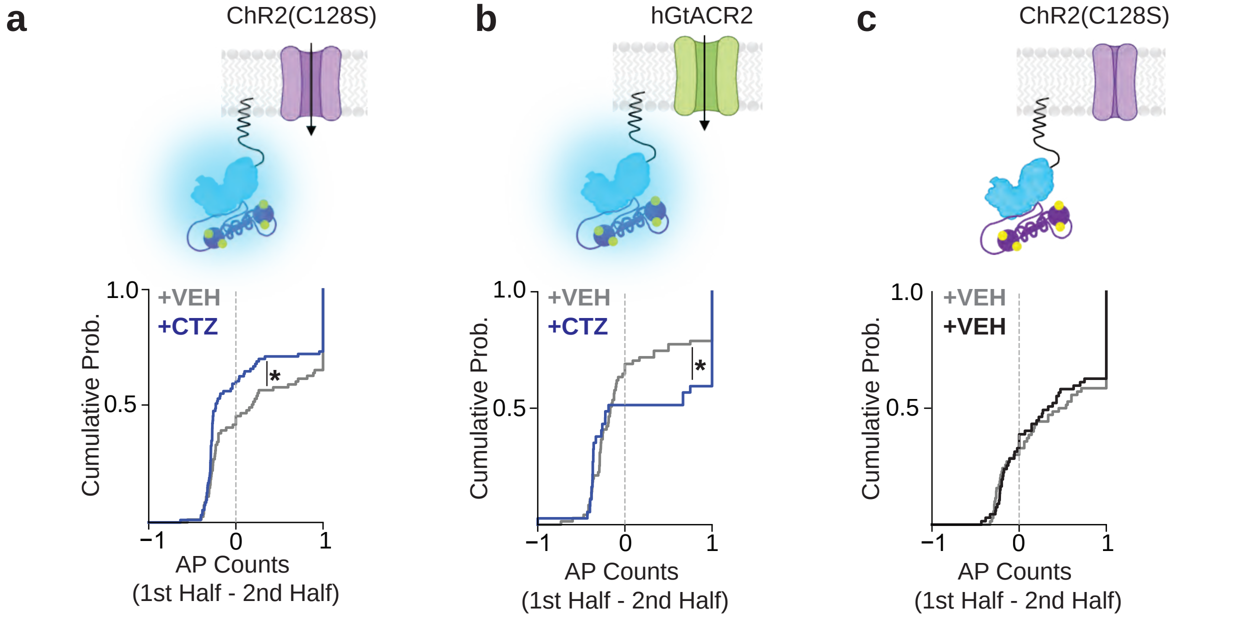


**Supplementary Figure S3. CTZ-dependent shifts in spike count distributions.** **a.** Excitatory opsin and CTZ perfusion. Cumulative distributions of the difference in action potential (AP) counts between the first and second halves of depolarizing current injections for neurons co-expressing LMC and the excitatory opsin ChR2(C128S) under CTZ perfusion. Gray traces correspond to recordings obtained during the initial vehicle perfusion, while blue traces correspond to recordings obtained during the subsequent CTZ perfusion. A significant rightward shift toward positive fraction-difference values was observed in the presence of CTZ (N = 16 neurons; Kolmogorov–Smirnov test, KS = 0.25, P = 0.01319), indicating enhanced firing in the second half of the stimulus. **b.** Inhibitory opsin and CTZ perfusion. As in (a), gray traces represent recordings during vehicle perfusion and blue traces represent recordings during CTZ perfusion. For neurons co-expressing LMC and the inhibitory opsin hGtACR2 under CTZ perfusion, cumulative distributions revealed a significant leftward shift toward negative fraction-difference values, reflecting reduced spike counts in the second half of the depolarizing current (N = 16 neurons; Kolmogorov–Smirnov test, KS = 0.3888, P = 0.00809). **c.** Excitatory opsin and vehicle control. Gray traces denote the cumulative distribution from the first vehicle perfusion, and black traces denote the cumulative distribution from the second vehicle perfusion. Neurons co-expressing LMC and ChR2(C128S) that received two sequential vehicle perfusions rather than CTZ during the second perfusion, showed no change in cumulative distributions and were indistinguishable (N = 14 neurons; Kolmogorov–Smirnov test, KS = 0.0597, P = 0.999).

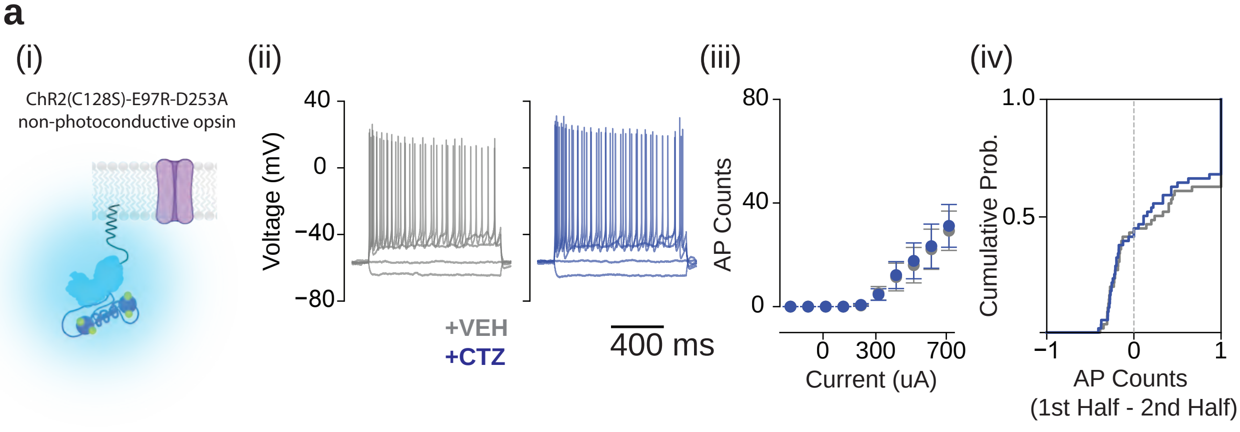


**Supplementary Figure S4. Lack of BLADe-mediated changes in neurons expressing a non-functional “DUD” opsin.** (i) Schematic of farnesylated LMC anchored to the inner cell membrane near the non-functional DUD opsin ChR2(C128S)-E97R-D253A showing that in the presence of depolarization-induced $Ca^{2}$ flux and luciferin (CTZ), no light-mediated modulation is expected. (ii) Representative membrane voltage responses to step current injections before (gray) and after (blue) bath perfusion of luciferin in a neuron co-expressing the DUD opsin and LMC, showing no change in firing. (iii) Action potential counts under vehicle (gray) and CTZ (blue) conditions remain similar. (iv) Empirical cumulative distribution of spike fraction changes across voltage traces for vehicle and luciferin conditions, demonstrating no shift in spike timing (Kolmogorov-Smirnov test, KS= 0.0545, P= 0.999) or firing rate in the absence of a functional opsin. Data expressed as mean ± SEM from N=6 neurons.

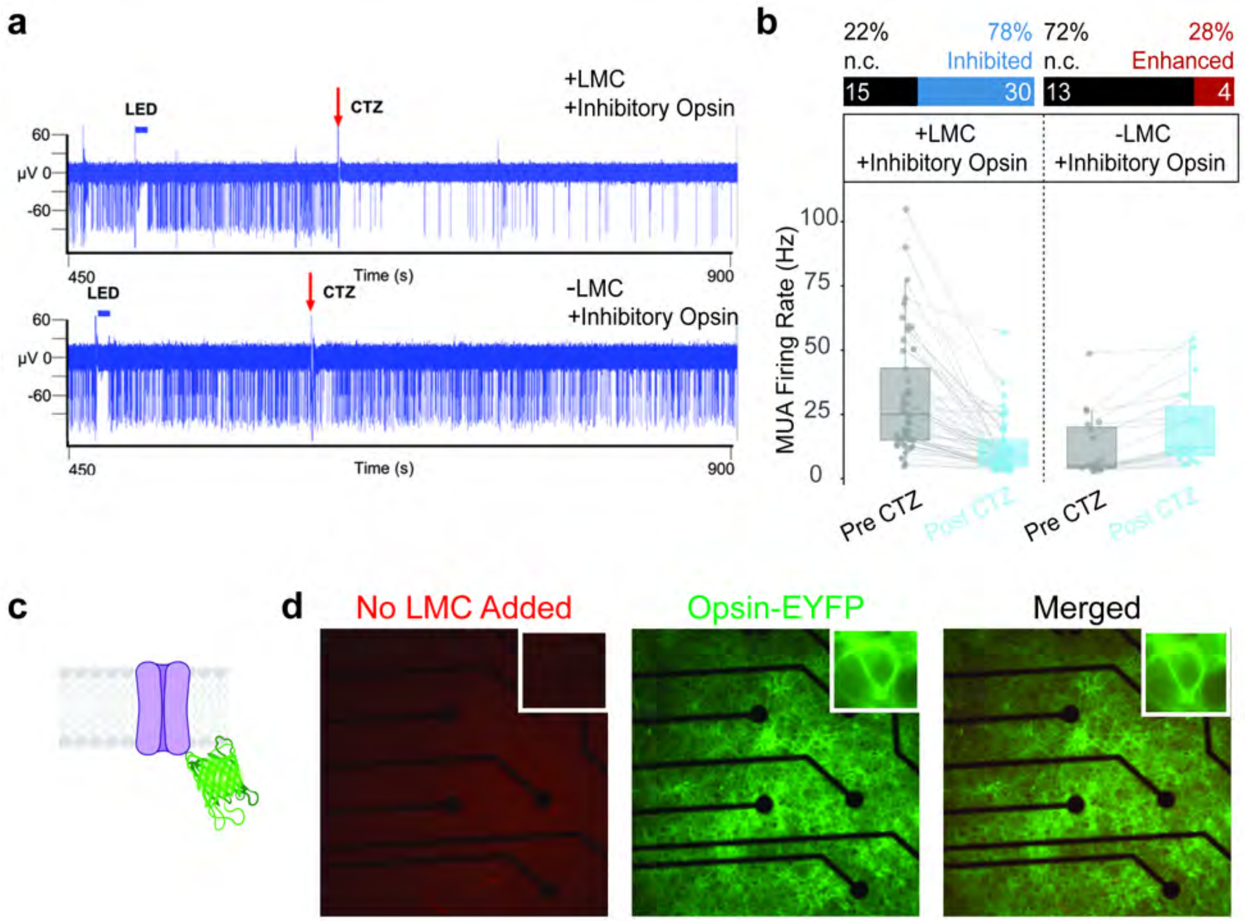


**Supplementary Figure S5. LMC enables light-dependent suppression of cortical network activity via an inhibitory opsin. a.** Representative multi-unit activity (MUA) traces from cortical neurons recorded on multielectrode arrays. Neurons co-expressing LMC and the inhibitory opsin hGtACR2 (top) show reduced spiking following luciferin (CTZ) addition (red arrow). In contrast, neurons expressing the opsin alone (bottom) show no consistent change in firing after CTZ. **b.** Quantification of firing rate changes before and after luciferin addition across individual electrodes. n.c., no change in firing; inhibited, decrease in firing; enhanced, increase in firing. **c.** Schematic of multi-electrode array (MEA) setup for opsin-only cultures. **d.** Fluorescence images of a representative opsin-only culture showing absence of dTomato signal (left, red channel) and strong opsin-EYFP expression (middle, green channel). Merged image (right) confirms that these neurons express the opsin but not LMC. Insets show magnified views of cell bodies overlaying electrode contacts. These cultures serve as a negative control for luciferin-dependent changes in network activity.

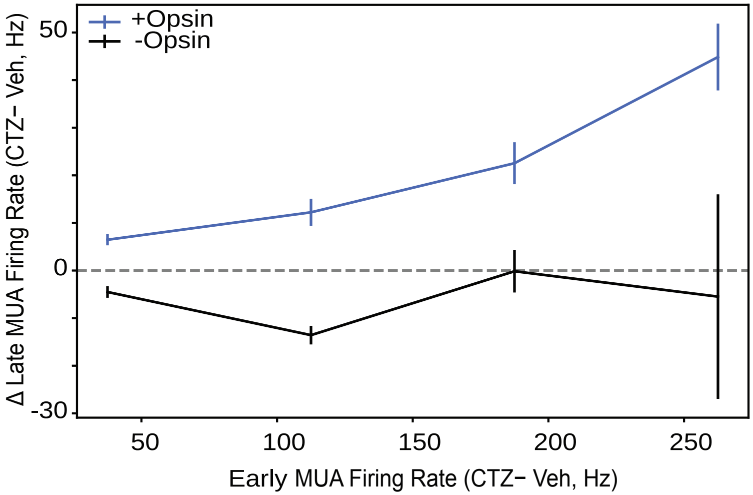


**Supplementary Figure S6. Activity-dependent modulation of late cortical activity by luciferin in vivo.** To visualize the relationship between early evoked activity and luciferin-dependent changes in late multiunit activity (MUA), we plotted the CTZ-induced change in late activity as a function of early evoked (0-200ms from stim onset) responses using binned trial averages. Early evoked activity was quantified shortly after stimulus onset (0-200ms from stim onset), and late activity (post 200ms from stim onset) was measured during a subsequent post-stimulus window. In opsin-expressing recordings (blue), the luciferin effect increased monotonically with early evoked activity, consistent with gain modulation. In contrast, recordings lacking opsin expression (black) showed no systematic dependence of luciferin effects on early activity. Error bars indicate confidence intervals.
